# Spatial and temporal control of lysis by the lambda holin

**DOI:** 10.1101/2023.09.08.556860

**Authors:** Jesse Cahill, Ashley Holt, Matthew Theodore, Russell Moreland, Chandler O’Leary, Cody Martin, Kelsey Bettridge, Jie Xiao, Ry Young

**Affiliations:** Sandia National Labs, Albuquerque, NM 87123; Center of Phage Technology, Department of Biochemistry & Biophysics, Texas A&M University, College Station, TX-77843, USA; Department of Biophysics and Biophysical Chemistry, Johns Hopkins University School of Medicine, Baltimore, MD 21205, USA

## Abstract

The infection cycle of phage λ terminates in lysis mediated by three types of lysis proteins, each disrupting a layer in the bacterial envelope: the S105 holin, the R endolysin, and the Rz/Rz1 spanin complex targeting the inner membrane (IM), cell wall or peptidoglycan (PG), and the outer membrane (OM), respectively. Video microscopy has shown that in most infections lysis occurs as a sudden, explosive event at a cell pole, such that the initial product is a less refractile ghost that retains rod-shaped morphology. Here, we investigate the molecular basis of polar lysis using time lapse fluorescence microscopy. The results indicate that the holin determines the morphology of lysis by suddenly forming two-dimensional rafts at the poles about 100 seconds prior to lysis. Given the physiological and biochemical similarities between the lambda holin and other class I holins, dynamic redistribution and sudden concentration may be common features of holins, probably reflecting the fitness advantage of all-or-nothing lysis regulation.

## INTRODUCTION

The *Caudovirales* (tailed phages) of Gram-negative hosts require three classes of lysis proteins directed to each component of the cell envelope [1]. The holin forms holes in the inner membrane (IM); the endolysin degrades the peptidoglycan (PG); and the spanins disrupt the outer membrane (OM) (Fig. 1A). In the λ infection cycle, these lysis proteins accumulate during the late or morphogenesis period beginning at about 8 min, resulting in lysis at 50 min, a time genetically programmed into the holin, S105 [1, 2]. S105, so named because it is a 105 aa product of the *S* gene, is the prototypical class I holin, an IM protein with three transmembrane domains (TMDs) disposed in an N-out, C-in topology (Fig. 1A) [3]. Prior to lysis, the S105 holins accumulate as dimers [4], freely mobile and uniformly distributed throughout the IM [5]. The timing of lysis is allele-specific to the *S* gene and varies dramatically with single missense changes, especially at positions within the TMDs. It has been proposed that lysis begins when the holin reaches a critical concentration that nucleates oligomerization into two-dimensional aggregates or “rafts” [5, 6]. Holin rafts have been visualized in studies with S105-GFP fusions [5]. In the current model for lysis, the rafts are thought to mediate a collapse in the membrane potential, which in turn promotes tertiary and quaternary rearrangement of the rafts into lethal lesions, or “S-holes” [1]. This aggregation/hole-formation process, unique to holins, had been termed “triggering” [3]. Cryo-electron microscopy and tomography studies revealed that the typical infection cycle results in micron-scale holes, typically 1 – 3 per cell and distributed randomly in the IM [5, 7, 8]. Experiments using non-invasive methods to assess the PMF revealed that that complete depolarization precedes lysis by ∼19 ±6 seconds [9]. Endolysin-mediated degradation of the PG requires holin function, presumably because the endolysin molecules, which accumulate in the cytoplasm with full muralytic activity, gain access to the PG when released through these holes into the periplasm. (Fig. 1Aii). Premature triggering can be caused by sudden loss of PMF, e.g. by energy poisons or an abrupt shift from aerobic to anaerobic growth [3, 10]. Titration with the uncoupler DNP revealed that triggering can be instigated with as little as a 30% decrease in the PMF [9].

**Figure 1.**
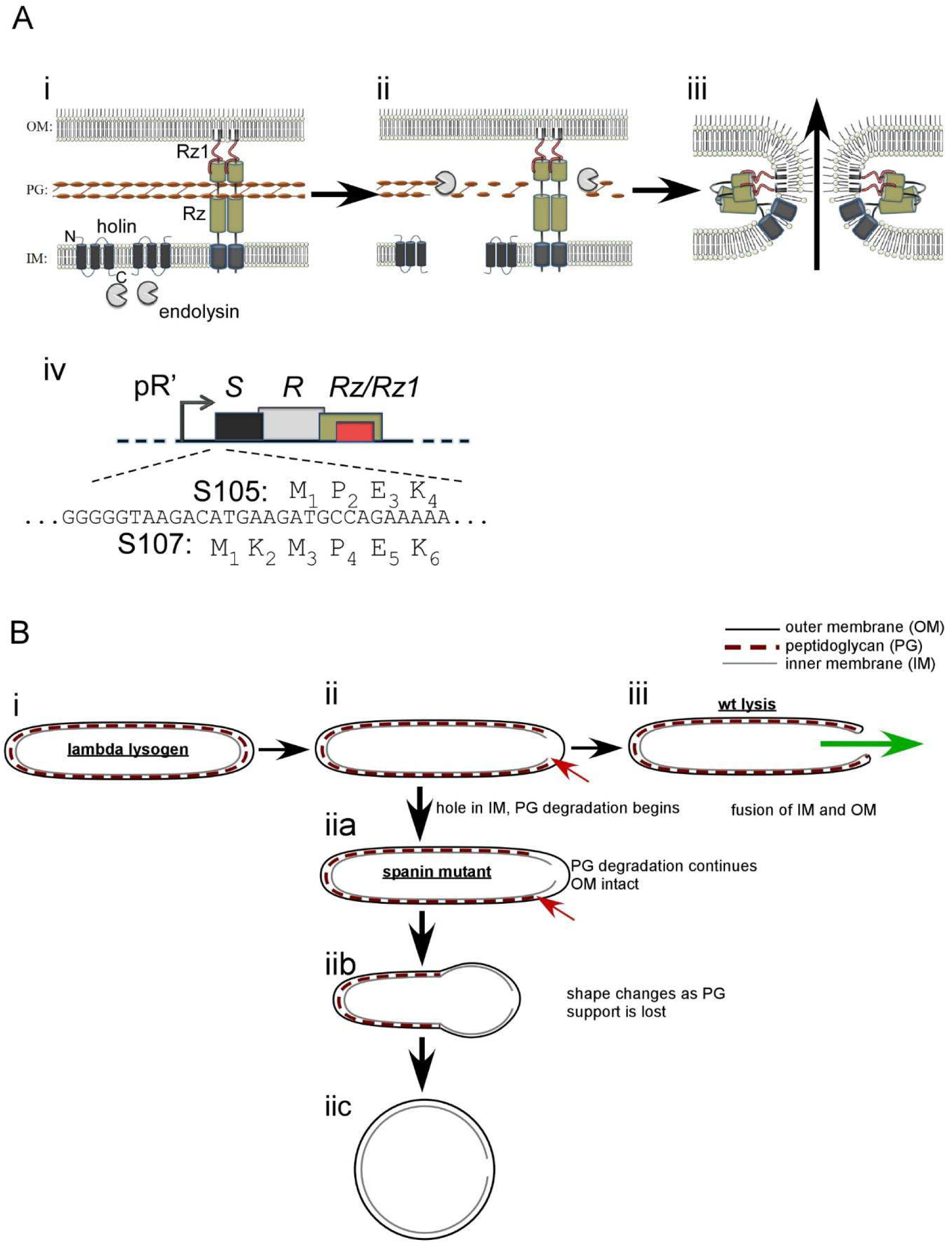
Overview of the λ lysis system. A) λ lysis genes and model for lysis protein function. (i) The topology and localization of lysis genes before lysis. (ii) The holin forms holes in the inner membrane (IM), which releases endolysin into the periplasm. Endolysin degrades peptidoglycan (PG). (iii) The spanin complex undergoes a conformational change that fuses the IM and outer membrane (OM), disrupting the OM. The arrow shows the direction of phage progeny release. (iv) The λ lysis cassette. The translational starts for the holin and antiholin (*S105* and *S107*) genes are shown above and below the DNA sequence, respectively. B) Model comparing the shape conversion of spanin mutants to lysis by wild-type λ. (i) λ lysogen before lysis. (ii) Hole formation in the IM by the holin and PG degradation is shown (red arrow). (iii) Lysis is complete after OM disruption. The putative fusion of the IM and OM is shown. The green arrow shows expected path of travel for intracellular content, including phage progeny. (iia) In the case of the spanin mutant, PG degradation continues (indicated by red arrow) without OM disruption. (iib) shape loss occurs in zone(s) where PG is absent. (iic) The terminal phenotype of spanin mutants is an un-lysed spherical cell.

After PG degradation, the last step of lysis in λ infections is destruction of the OM [11, 12]. This is accomplished by the spanin complex, composed of two subunits: Rz, an integral IM protein; and Rz1, an OM lipoprotein (Fig. 1A) [12, 13]. Each subunit forms a covalent homodimer, linked by intermolecular disulfide bonds. Each Rz homodimer forms a complex with an Rz1 homodimer via C-terminal interactions, resulting in a heterotetramer that is threaded through the meshwork of the PG to span the entire periplasm. Destruction of the PG is proposed to result in liberation of the spanin complexes from the encaging PG meshwork and thus activating them for completion of the lysis pathway (Fig. 1A ii-iii, B) [11]. In most cells, lysis is observed as explosive events in which the cellular contents escape from a single S-hole, resulting in the generation of non-refractile ghosts that at least temporarily retain rod shape [11].

However, in spanin mutants, the infected cell does not lyse but instead is transformed to a spherical shape once the PG is degraded. Cryo-EM studies revealed that the OM is intact in these spherical forms. (Fig. 1B ii a-c) [14]. The similarity of spanins to class-I viral fusion proteins, in terms of linkage of apposed membranes and the prominent coiled-coil domains, led to a model in which spanins disrupt the OM by fusing the IM and OM (Fig. 1A iii). Recent experiments with spheroplasts displaying the periplasmic domains of Rz and Rz1 confirmed that these proteins are fusogenic [15].

Here we investigate the basis of phenotypic changes that occur during the terminal lytic event real time. Using high resolution microscopy techniques, we monitor lysis morphology at the single cell level and determine that the site of sudden holin rearrangement prior to lysis is spatially correlated with where the lytic blowout occurs. The results are interpreted in a detailed model for the λ lytic pathway.

## METHODS

### Strains, bacteriophages, plasmids and growth conditions

The *Escherichia coli* K-12 derivative MG1655 *lacI^q^*Δ*lacY* was used as a host in this study. Bacteriophages, plasmids, and strains used are listed in Table 1. The use of selection markers within phages was shown to have no effect on wild type lysis [14]. Overnight cultures were made in LB supplemented with 100 µg/mL ampicillin, 40 µg/mL kanamycin, or 10 µg/mL chloramphenicol when appropriate. Growth of cultures and lysis profiles were monitored as described previously [13]. Briefly, overnight cultures were diluted 1:200 in 25 mL LB in a 250 mL flask supplemented with the antibiotics above and 10 mM MgCl_2_ unless otherwise indicated. Lysogens were incubated at 30 °C and aerated at 250 RPM. At A550 ∼0.3, lysogens were induced by aerating at 42 °C for 15 min and continued growth at 37 °C until lysis. When indicated, isopropyl β-D-thiogalactopyranoside (IPTG) was added at the time of induction at a final concentration of 1 mM. For induction of *phoA-R* under P_araBAD_ control (pSec-R), arabinose was added at a final concentration of 0.4% at 25 min after induction.

**Table 1.**
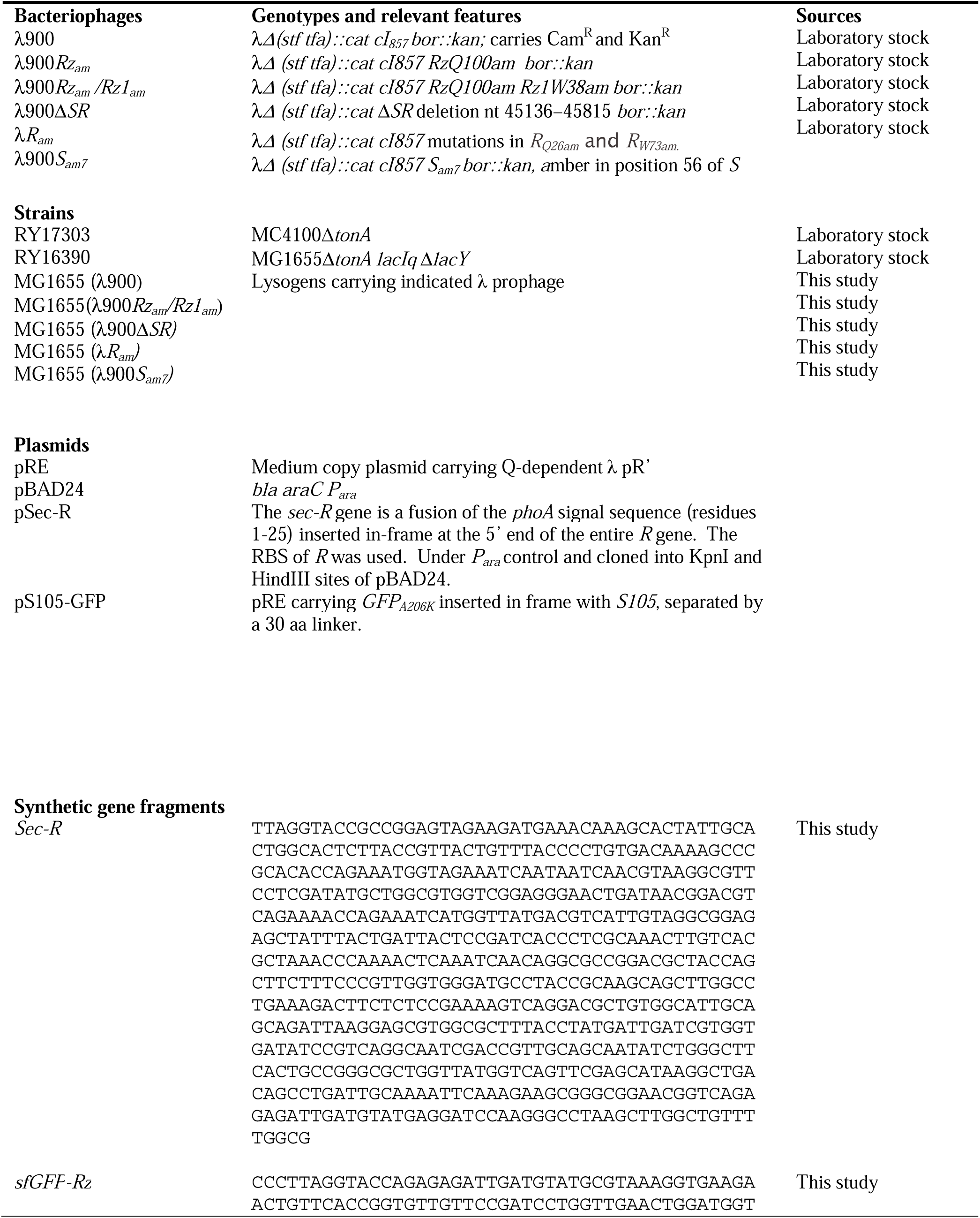
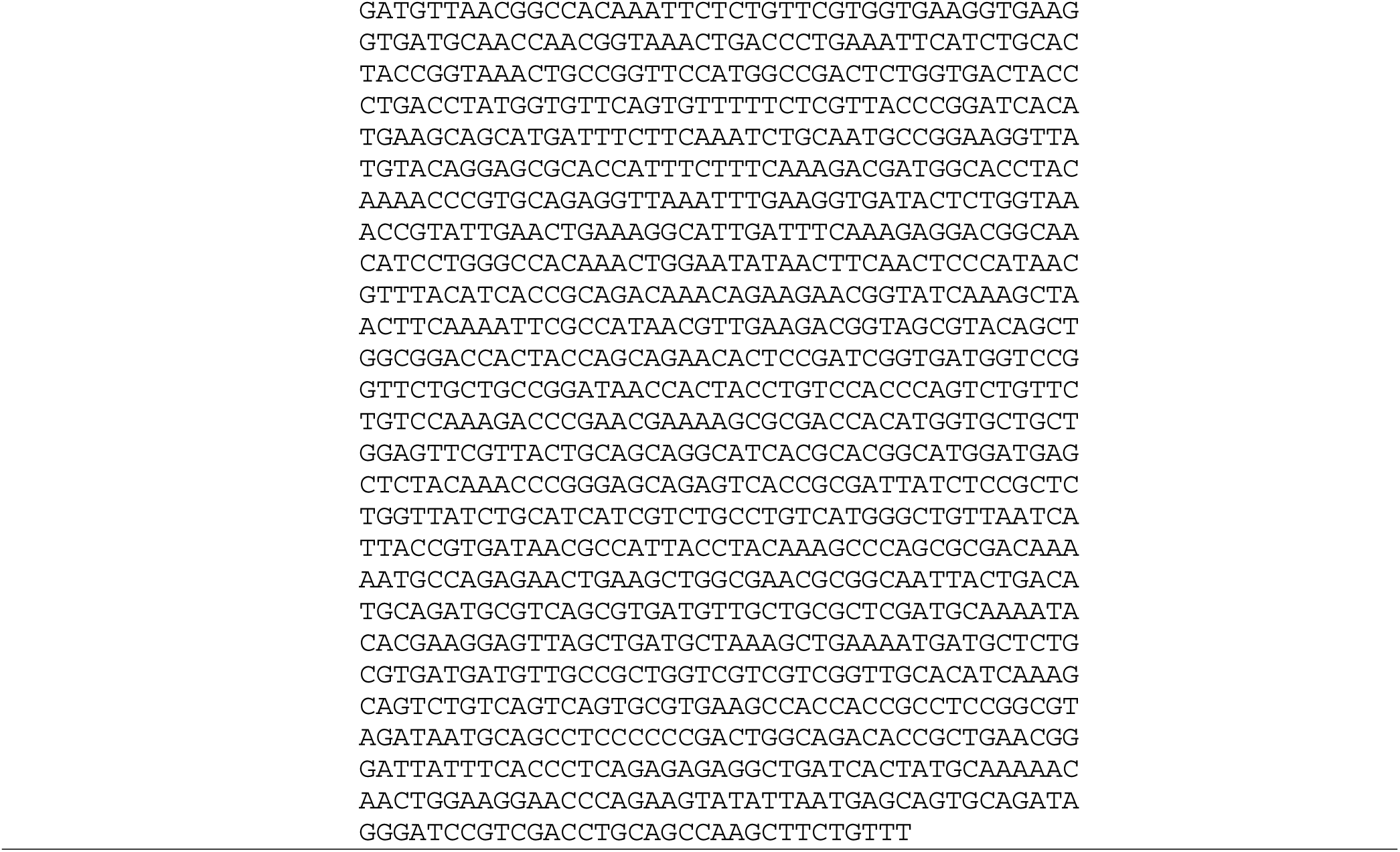
Phages, strains, plasmids, and synthetic gene fragments used in this study.

### DNA manipulation

Plasmid DNA isolation, PCR, amplification, site-directed mutagenesis, transformation, subcloning, and sequencing were performed as described previously [13]. Synthetic gene fragments (gBlocks) were obtained from IDT (Integrated DNA Technology, Coralville, IA).

Restriction modification was performed using enzymes purchased from NEB (New England Biolabs) according to manufacturer’s specifications. Primers 5’- ACGATGTGC<u>AT</u>CATTATCGCCTGGTTCATTCG-3’ and 5’- GCGATAATG<u>AT</u>GCACATCGTTGCGTCGATTACTG-3’ (changed nucleotides are underlined) were used to introduce A52I substitution into the *S105* and *S105-GFP* genes.

### Phase contrast time-lapse microscopy

All samples were collected for imaging at 45 min after induction except for Ethylenediaminetetraacetic acid (EDTA) experiments. Subcultures used for monitoring lysis in the presence of EDTA were not supplemented with MgCl_2_. At 42 min after induction, five mL of 0.5 M EDTA were added to 20 mL of aerating cultures. Shaking was continued for five min and cells were imaged immediately. All samples were prepared and monitored the following way: A 1.5 µL aliquot of sample was placed on a glass slide and covered with a coverslip. Cells were imaged immediately using a plan-apochromat 20x/0.8 Ph2 objective installed on a Zeiss Axio Observer 7 inverted microscope. Time-lapse video was captured at 0.5-5 frames per second for 10 min or until lysis or shape change was complete. When cells were in different focal planes, Z-stacks were included in the time-lapse program to maintain focus on all cells. To maintain a stable temperature of 37 °C on the mounted slide, a Lab-TekTM S1 heated insert was pre-heated to 42 °C, and temperature control was active for the duration of the experiment.

Video scaling, editing, measurement, and image export was done using the Zen 2.3 software. The subcellular site of blebbing and lysis were scored based on observed morphological changes by dividing the long axis of cells into five equal compartments. Using this convention, cells have two polar compartments, and three compartments within the sidewall of the cell (two parapolar and one midcell) (See Fig. S1).

### Fluorescence microscopy

Samples were collected and prepared for imaging as described above, except cells expressing *S105-GFP* were collected at 49 min after induction. For experiments monitoring Thioflavin-T (ThT) fluorescence, ThT was added from a 1000x stock at the time of thermal induction at a final concentration of 10 µM. ThT was purchased from Sigma-Aldrich (St. Louis, MO); stocks were made in filter-sterilized water and kept at -20 °C prior to use. ThT time-lapses were captured at 2 frames per second using Zeiss filter cube 91 HE CFP illuminating at 401-445 nm with 20 ms exposure time at 50% light intensity. GFP time-lapses were captured at 1-3 frames per second using a Zeiss filter cube 90 HE (DAPI/GFP/Cy3/Cy5 multi-band pass filter cube) illuminating at 450-488 nm with 40 ms exposure time at 50% light source intensity using a 100x/1.46 Ph3 objective. Brightness and contrast were adjusted with Zen 2.3 software setting the area outside of cells to be the background. Raft number and location was determined by foci apparent in the GFP channel. Rafts were determined to be polar, parapolar, or midcell-localized by dividing the long axis of the cell into five equal compartments (described above, and shown in Fig. S1).

### Super-resolution cell preparation

Cells were collected at 50 min after induction and fixed in fixation solution (2.6% paraformaldehyde and 0.8% glutaraldehyde in 1x PBS solution). Fresh paraformaldehyde and glutaraldehyde were used in each experiment (Electron Microscopy Sciences #15710 and #16019), and 10x stock of PBS was used to create 1x PBS solution (Quality Biological #119- 069-101). Cells were fixed at room temperature for 15 min, then washed twice in 1x PBS solution. Cells were permeabilized in 1x PBS with 0.1% Triton X-100 for 5 min at room temperature without rotation, then washed twice in 1x PBS solution. The cell wall was minimally degraded by treatment in 1x PBS with 10 µg/mL lysozyme for 5 min at room temperature without rotation, then washed twice in 1x PBS solution. The presence of lysozyme was required for the antibody to permeate the cell wall (data not shown). Cells were next incubated in 1x PBS with 10% goat serum (Sigma-Aldrich #G9023) at 30 °C for 30 min with rotation. The S105 antibody [16] was added in a 1:100 dilution directly to this mixture, and cells with antibody were incubate at 4 °C overnight (16-18 hours). Cells were washed once in 1x PBS with 0.05% Tween- 20 (PBST solution) and then incubated in 1x PBS with a 1:250 dilution of GAR-AF647 (Thermo Fisher Scientific #A32733) secondary antibody for two hours at room temperature with rotation. Cells were washed with PBST solution four times and then resuspended in a small volume of 1x PBS solution and stored at 4 °C until ready to image. Cells were adhered to coverslips treated with 0.1% poly-L-lysine and bathed in STORM buffer. STORM buffer consisted of 10% glucose w/v, 40 mM cysteamine (Sigma-Aldrich #30070), 40 ug/mL glucose oxidase (Sigma-Aldrich #G7141), 3 ug/mL catalase (Sigma-Aldrich #C3515) in a 1x TBS solution.

### Super-resolution STORM imaging and analysis

Immobilized cells were imaged on an Olympus IX71 inverted microscope with a 1003, 1.49 NA oil-immersion objective under widefield illumination. To excite fluorophores, we used solid-state lasers with wavelengths at 405 nm and 647 nm (Coherent OBIS lasers) set to 1 mW and 100 mW respectively (approximately 3.5 W/cm^2^ and 2.9 kW/cm^2^ respectively). Images were collected using MetaMorph software (Molecular Devices); emitted photons were collected using an EMCCD camera (iXon Ultra 897, Andor Technology) under an EM gain of 300. Exposure time of each frame was kept constant at 10 ms per frame, with movies lasting a total of 3000 frames. Between 12 and 15 movies were taken of each region for a total of 36,000 to 45,000 frames per region. The region imaged was 150 by 150 pixels (160 nm per pixel), and an emission filter (Chroma ET700/75m) was used to reduce background. Brightfield images of cells were taken between movies to correct for any x-y drift. Movies were imported into ImageJ, and substacks were taken (frames 2601-3000 were used for the first movie, and 501-3000 were used for subsequent movies). Movies were concatenated together to form the “STORM sequence” and saved as a tiff stack. The STORM sequence tiff stack was analyzed using the ThunderSTORM plugin (Wavelet filter (B-spline) with scale 2.0, order 3, using the local maximum detection at 2 times the standard deviation of the Wavelet filter, and fitting of spots used a Gaussian PSF with a starting sigma of 1.5 pixels and starting radius of 3 pixels using a maximum likelihood algorithm). Resulting coordinates were imported into MATLAB for further post-analysis measures. Images were filtered based on sigma and uncertainty in fitting, and in-house software was used to correct for blinking. Typically, between 30,000 and 80,000 localizations were obtained per region after these corrections. Images were created using in-house MATLAB software using a sum-of-Gaussians approach, with the size of the molecule in the resulting image related to the uncertainty in fitting of the molecule. Resolution was determined through the thunderSTORM plugin, and averaged around 35 nm.

### Structured Illumination Microscopy (SIM) and analysis

Immobilized cells were prepared as above (Materials and Methods, subsection “Super-resolution cell preparation”) with the exception that cells were incubated with GAR-AF488 antibody (Thermo Fisher Scientific # A-11008) and were bathed in anti-fading buffer (1x PBS, 60% glycerol, 0.2% propyl gallate (Sigma-Aldrich P3130)) instead of STORM buffer. Cells were imaged using a Deltavision OMX-SR (General Electric) microscope using 20 ms exposure and performing z-stacks on ∼3 μm of z-space (each z-stack has a resolution of 125 nm, so 3 μm of z-space equates to 24 z-stacks). The 488 nm excitation laser was used at 0.5% of the maximum power. GE software softWoRx v7.0 was used to reconstruct the images to create the three-dimensional z-stack of cells. Reconstructed images were then imported into Fiji and saved as tiff stacks. Images were visualized using the “gem” look-up table in Fiji. To determine the percent of cells in which mid-plane rafts were detected, images were manually inspected to locate the rafts on the surface of the cell. Cell were considered to have mid-plane rafts if fluorescence signal could be continually localized to the same region from the membrane to the mid-cell.

### Western blotting

Protein samples were collected as described previously for Rz [17]. Briefly, a 1 mL aliquot of whole-cell sample was taken at 45 min after induction. Samples were precipitated by tricholoroacetic acid (TCA), and pellets were resuspended in sample loading buffer normalized to the A_550_ units at the time of collection. After 5 min of boiling, 0.3 units of sample were resolved on a Novex Wedgewell 4-to-20% Tris-Glycine SDS-PAGE gel (Thermo Fisher). Gel transfer and immunodetection were done using the iBlot and iBind systems (Thermo Fisher) according to the manufacturer’s recommendations.

## RESULTS

### Lysis polarity is lost with host-secreted endolysin

Previously, the lysis morphology of induced λ lysogens was monitored by phase time-lapse microscopy, revealing that 34 of 40 cells ruptured from a pole [11]. Moreover, in inductions of spanin-defective lysogens, where the terminal phenotype was spherical cells, the transformation from rod to sphere began with deformation at the poles in 39 of 44 cells. These results indicated that in holin-endolysin mediated lysis, degradation of the PG usually begins at the poles. Theoretically, inhomogeneity in the susceptibility of PG to muralytic enzymes leaves the simple possibility that polar lysis could simply reflect greater sensitivity to the endolysin at the poles. To test this notion, we compared the morphological changes in situations where the endolysin was either released to the periplasm by S105 hole-formation or secreted to the periplasm via a normal signal in the absence of holin function. As before, a large majority of induced cells underwent polar lytic events in S105-mediated lysis (Figure 2A, Movie 1) and, in the spanin-negative condition, began rod to sphere transformation at the poles (Fig. 2B, Movie 2). However, with the secreted endolysin, the induced cells converted from rod to spherical shape isotropically; i.e., the long axis contracted while the short axis expanded uniformly (Fig. 2C, Movie 3). This suggests that the endolysin degrades PG in a uniform manner. In other words,the mode of delivery (i.e. S-holes) determines the polar localization. Thus neither spanin function nor preferential degradation of the polar PG is necessary for polar lysis.

**Figure 2.**
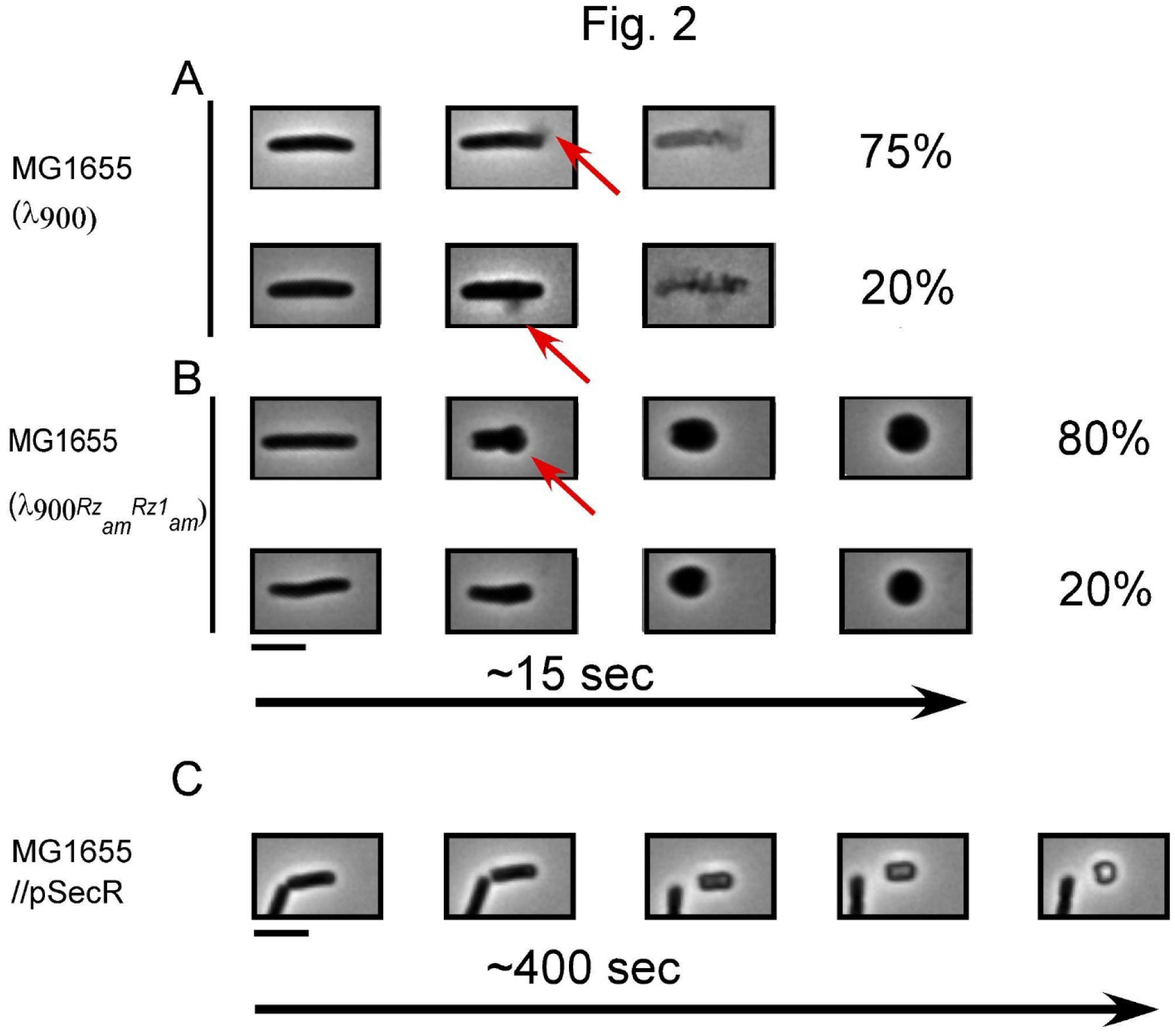
Lysis morphology and shape conversion of λ lysogens, spanin mutants and cells expressing secreted endolysin. MG1655 lysogens were thermally induced and monitored by phase contrast microscopy 1 min prior to lysis. The average time from the first frame prior to an observable morphological change to lysis or the completion of rod-to-sphere shape conversion is displayed below the micrographs. Micrographs are representative of the lysis morphology or shape conversion observed. Percentages are given to the right of micrographs to indicate how frequent the displayed type of morphological change occurs. A) λ900 lysogens showed two primary sites of rupture (indicated with red arrows): in the side wall and in the polar region. The remaining 5% in panel A are instances when the site of rupture was unclear. B) Similar to the lysogens in panel A, λ900*Rz_am_Rz1_am_* lysogens showed shape conversion beginning either at the poles or by apolar inflation. C) MG1655 cells expressing p*Sec-R* were induced with 0.4% arabinose and monitored as described above. n-values are 134, 40, and 62 for A, B, and C, respectively. The red arrows are marking the site of lytic blowout or shape conversion. Scale bar = 5 µm.

### Blebbing in EDTA-treated cells reveals sites of hole formation

EDTA reduces the strength of the OM by sequestering divalent cations [18]. We hypothesized that in the absence of spanins, the sudden formation of the micron-scale holes in the IM would result in bleb formation in the juxtaposed OM after EDTA-treatment (Fig. 3A). To test this notion, EDTA was added to the induced λ900*Rz_am_Rz1_am_*culture five min prior to the time when rod-to-sphere conversion occurs. After incubation with EDTA, an aliquot of cells was withdrawn, and fields were randomly selected and monitored using phase time-lapse microscopy.

**Figure 3.**
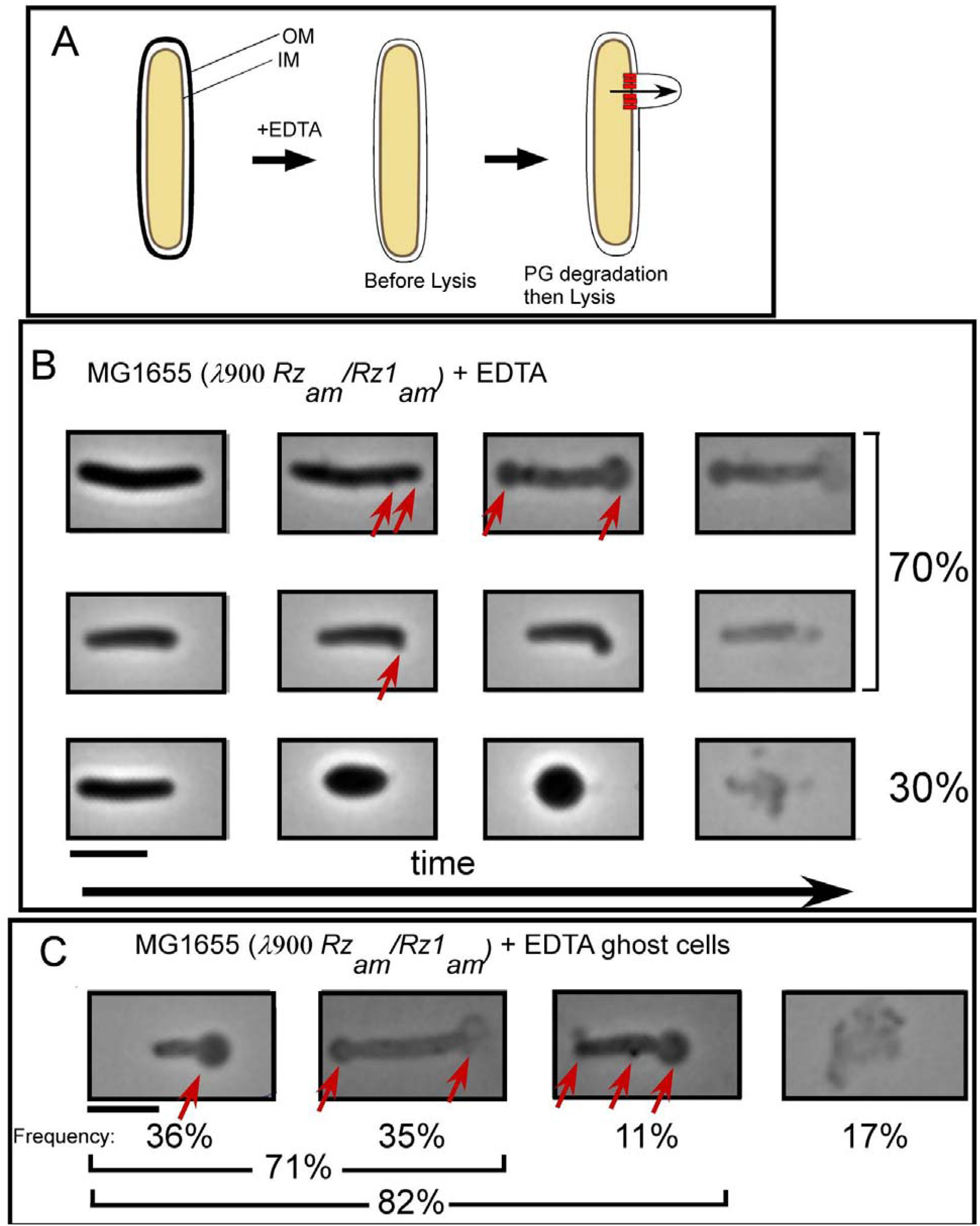
EDTA complements the lysis defect of spanin mutant cells. A) Cartoon of morphological changes expected upon EDTA treatment of spanin mutant lysogens. B) Representative time-lapse series of λ900*Rz_am_/Rz1_am_* lysogens treated with EDTA 5 min prior to lysis. Micrographs are representative of the lysis morphology or shape conversion observed. C) Ghost cells of spanin mutant lysogens treated with EDTA. Red arrows indicate blebs or blowouts. Percentages are given to the right of micrographs indicate how frequent the displayed type of morphological change occurs. The four major representative classes of ghost are shown. n = 122 ghost cells. Scale bar = 5 µm.

Cell morphology remained predominantly rod-shaped, similar to wt lysogens in normal conditions (Table S1). Out of 44 cells, 70% formed blebs at the poles before lysing (Fig. 3B, Table S1, Movie 4 and 5). Some cells (n=15) showed bleb formation in the sidewall of the cell along with polar blebs. Sidewall blebs were smaller, and the deformation was local and did not change the overall rod shape of the cell. There were also cells (n=13) that converted to spherical shape before lysing, as observed with cells that are not treated with EDTA (Movie 6 and compare the third set in Fig. 3B to Fig. 2B). Importantly, these cells lost rod shape uniformly, without deforming from either pole. The simplest explanation is that the endolysin is released to multiple areas of the sacculus at the same time. In other words, we suspect that is the result of a polydisperse distribution of S-holes.

After lysis, cells are recognizable as rod shapes with reduced refractility. Such cells are commonly referred to as “ghosts,” and the morphology of cell ghosts have been shown to correlate with lysis morphology [11]. Therefore, we analyzed 122 ghosts of λ900*Rz_am_/Rz1_am_*lysogens treated with EDTA. Morphologically, all ghosts showed signs of having formed at least one polar bleb and most ghosts (82%) were rod-shaped (Fig. 3C). Most ghosts (71%) had 1-2 apparent polar blebs and 11% of ghosts appeared to have formed a bleb in the sidewall in addition to a polar bleb. In both time-lapse and ghost analyses of λ900*Rz_am_/Rz1_am_*lysogens treated with EDTA, the percentage of polar blowouts was consistent with the percentage of polar lysis of wt lysogens without EDTA treatment. Taken together, it is clear that EDTA-treated lysogens expressing only the holin and endolysin predominantly form large blebs at the poles prior to lysis, supporting the notion that it is the holin that is required for the bias towards polar lysis.

### Raft formation by a holin-GFP fusion

Previous studies with S105-GFP allowed visualization of holin raft formation in inductions in a background where lysis was abolished by ablation of the endolysin and spanin genes [5]. Here we again exploited this chimera in studies with fully functional endolysin and spanin genes to focus on the relationship between raft formation and lysis, using a much higher frame rate (2 sec vs 1 min) [5]. We monitored lysis of λ900*S_am7_* lysogens in which the prophage holin defect was complemented by *S105-GFP* (Fig. 4A) expressed from a medium copy plasmid under native late gene transcriptional control, conditions which have previously been shown to recapitulate normal lysis kinetics. [5, 19, 20].

**Figure 4.**
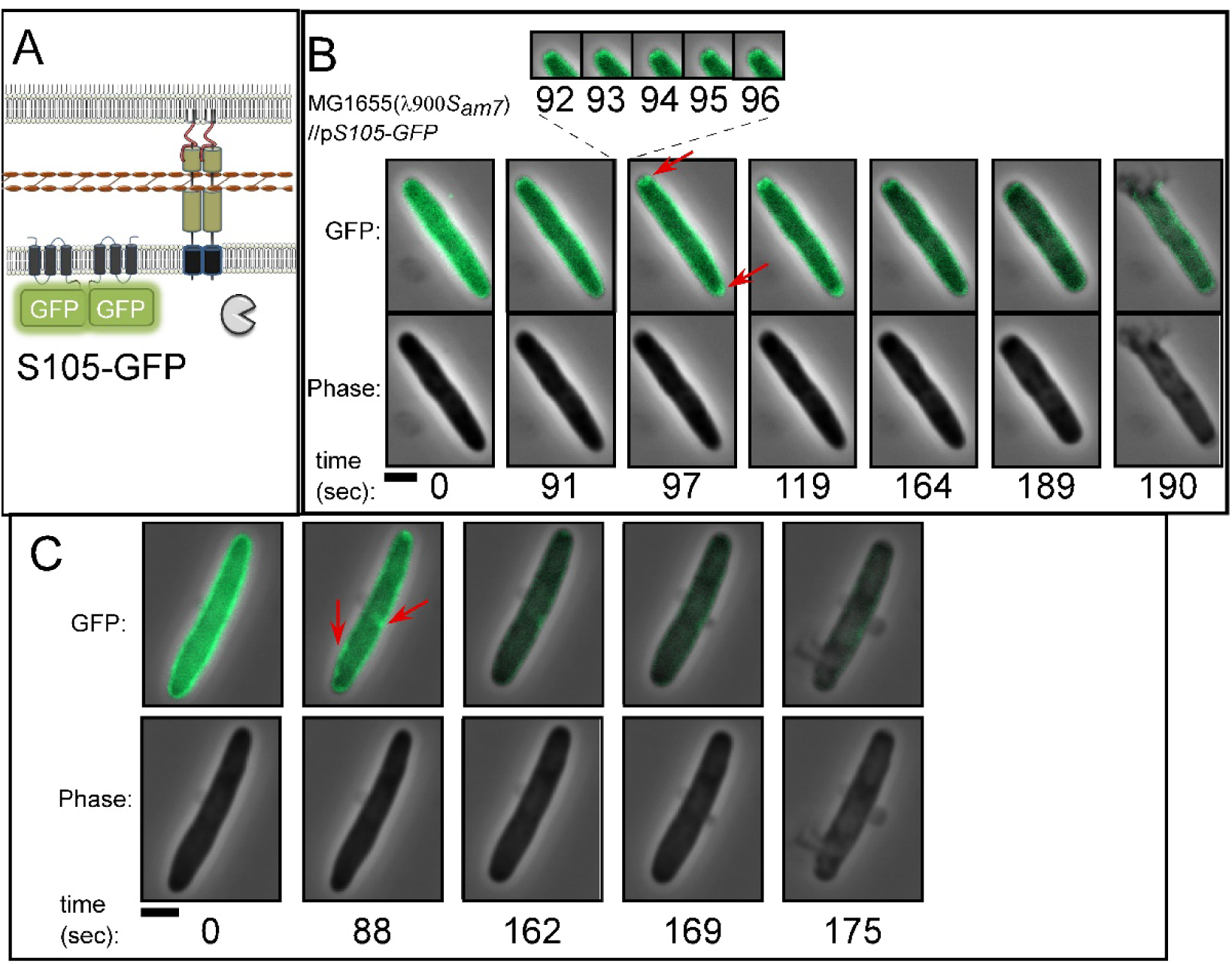
Lysis of λ900S*_am7_* lysogens expressing p*S105-GFP*. A) Cartoon showing the localization of lysis proteins produced by λ900S*_am7_* plasmid- borne p*S105-GFP* B) and C) Representative time lapse images of isogenic λ900S*_am7_* lysogens induced for *S105-GFP*. Cells were imaged 1 min before lysis. Time is shown below panels. Red arrows indicate rafts. Scale bar = 2 µm

Fifty single cell lysis events from induced cultures were monitored by time-lapse fluorescence microscopy (Table S2). Overall, the polar bias was retained by the fusion allele, although the degree was marginally decreased (56% of cells). In most (39/50) cells, the S105- GFP signal was uniformly distributed at the onset of monitoring. At an average of 93 sec (± 22) before lysis, the S105-GFP signal suddenly formed foci, or rafts (Fig. 4B, Movie 7, Table S2). On average, we detected 2.3 rafts per cell (n=46), and rafts were most often associated with the poles (Table S2). Just prior to lysis, the raft dissipated in most cells (n=41) on average ∼35 ± 20 sec (n=41) before lytic blowout (Table S2). In most cases (65%), raft formation was correlated to the site of lysis, even when lysis did not occur from the poles (Fig. 4C). Moreover, the formation of rafts was sudden, occurring within 2-3 seconds (Fig. 4B). Interestingly, rafts were sometimes unstable, forming foci, then delocalizing before forming foci again (Fig. 5, Movie 8) (n=6 cells out of 46). In Table S2, we refer to raft instability as “flickering.” In one such case, raft formation was completely abortive: a raft was initially detected at an area of curvature within the peripolar region. This raft dissolved, and then new rafts formed at a different site (the poles) (Fig. 5B). Taken together these data demonstrate cell-to-cell differences in the way the holin rearranges prior to lysis and suggest that the distribution of the holin rafts is likely the primary determinant of lysis morphology (discussed later).

**Figure 5.**
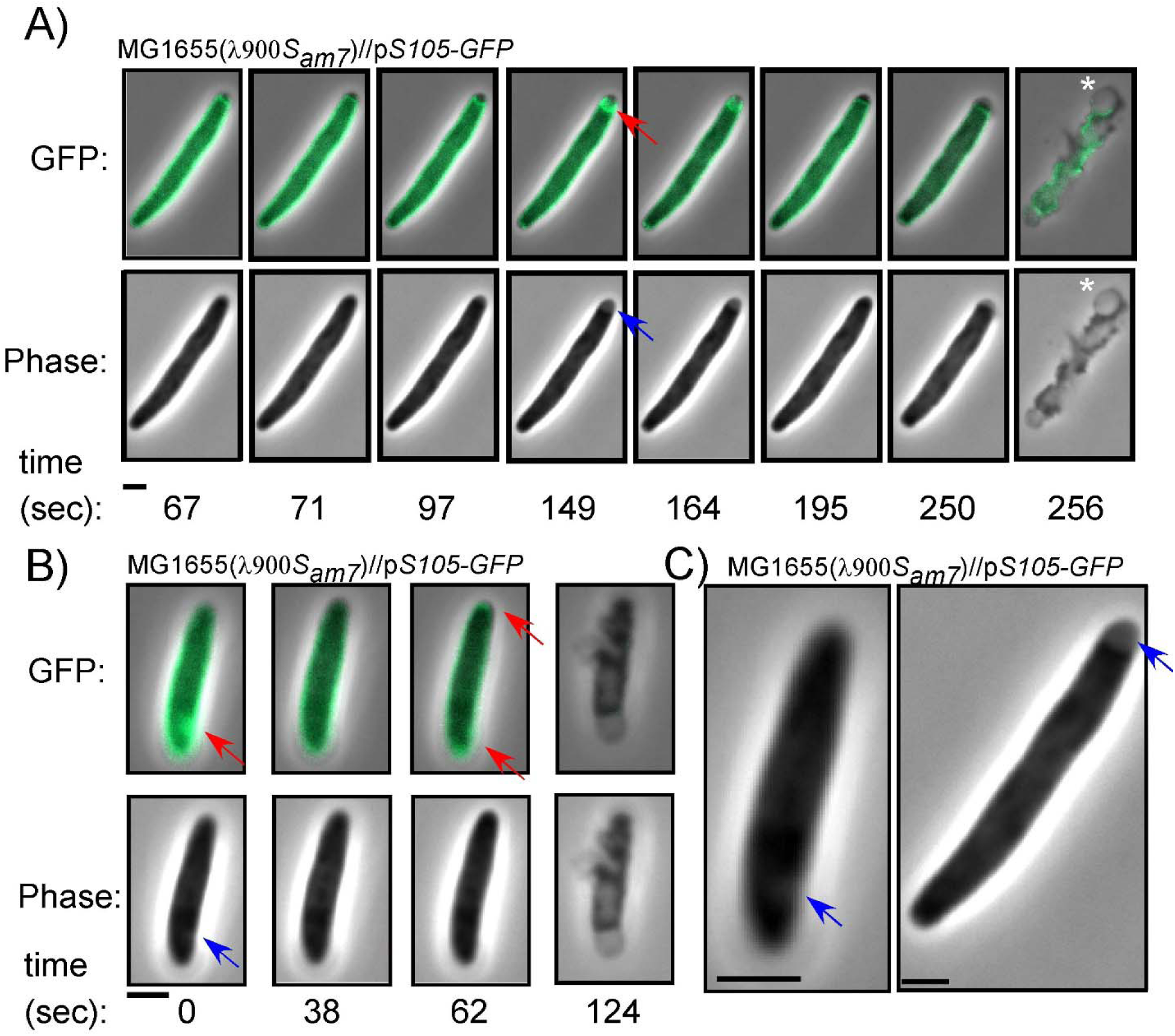
The dynamics and features of S105-GFP rafts. λ900S*_am7_* lysogens induced for lysis and expressing *S105-GFP*, isogenic to Fig. 4. A and B) Two examples of intermittent holin raft formation (denoted “flickering” in the text) can be seen. Arrows indicate rafts (red) and phase-light spots or plasmolysis bays (blue). Vesicles generated after lysis are denoted by asterisks. C) inset of cells showing plasmolysis bays. Scale bar = 2 µm.

Out of the 46 cells that formed rafts during these experiments, most (33 cells) had phase-light spots associated with at least one raft (Table S2). These spots were concurrent with raft nucleation and were morphologically similar to plasmolysis bays, where the inner membrane retracts from the rest of the cell envelope [21] (Fig. 5A-C, blue arrow). Although the basis for this is unclear, it suggests that the formation of rafts in some cases leads to an invagination of the IM.

### S105 A52I mutation produces polar rafts that do not form holes

The data above suggest that the site of raft formation determines the site of lysis. Therefore, in order to better characterize the superstructure and position of rafts, we sought to identify an *S105* mutant that is blocked at the rafting step. Previously, substitutions at position A52 within the second TMD helix were shown to have an extreme and unpredictable effect on holin function [22–24]. For example, the A52V allele fails to trigger and is blocked before raft formation, whereas substitutions with Gyl or Leu result in severe early triggering [5, 25]. In a saturating in vitro mutagenesis study it was discovered that an Ile substitution is non-lytic with an even tighter lysis-defective phenotype than A52V [25]. Based on this, we wondered whether non-lytic *S105_A52I_*would be defective for formation of rafts like A52V or stalled at a later step in holin function. As was shown with *S105_A52I_* [25], the A52I change also blocked holin function in the S105-GFP background (Fig. 6A), and western blotting indicated that this product accumulates normally (Fig. 6B). When imaged at the time of lysis, most lysogens expressing *S105_A52I_-GFP* had formed rafts (n = 235/260) (Fig. 6C and D), which was in contrast to the persistent uniform peripheral signal previously shown to be generated by the A52V product [5]. Like wt S105-GFP, the pattern of S105_A52I_-GFP signal prior to triggering was uniform and peripheral (not shown). Stable rafts formed ∼60 min after induction. The raft foci were visible in the phase contrast channel and were similar in appearance to plasmolysis bays [21] (Fig. 6D, blue arrow). We took z-stacks to investigate our previous interpretation that phase-light spots were a result of rafts causing an invagination of the IM. Although an apparent ring-like structure could be observed in some z-stacks (Fig. 6D, red arrow), deeper sections revealed that the GFP signal of the A52I rafts were instead cup-shaped (Fig. 6D, stacks -240 and -480). This is consistent with the notion that the rafts formed by S105_A52I_-GFP create a depression on the membrane. Although the molecular basis for the defect is unknown, A52I rafts fail to progress toward a hole-formation. Notably, the rafts produced by *S105_A52I_-GFP* were localized to the poles 82% of the time (213 out of the 235 cells with rafts). In contrast to S105, the A52I allele produced ∼1 raft per cell on average (223 cells with single rafts/235 cells). To assess whether these data can be recapitulated within the native S105 context, we performed Immuno-STORM imaging of λ*R_am_* lysogens. Cells were fixed when the wild-type λ strains begin to lyse (t = 50 min post thermal induction) to determine their distributions at this high concentration of holin (Fig. S2A). We found that overall, the distributions were similar to that of S105-GFP images, suggesting fixation does not drastically alter the distribution of the holin molecules. The distribution of holin appears to be somewhat homogenous across the membrane, with some moderate preference for localizing to the poles, consistent with above results. On average, the cells had 2.67 ± 0.83 rafts per cell, consistent with the S105-GFP images.

**Figure 6.**
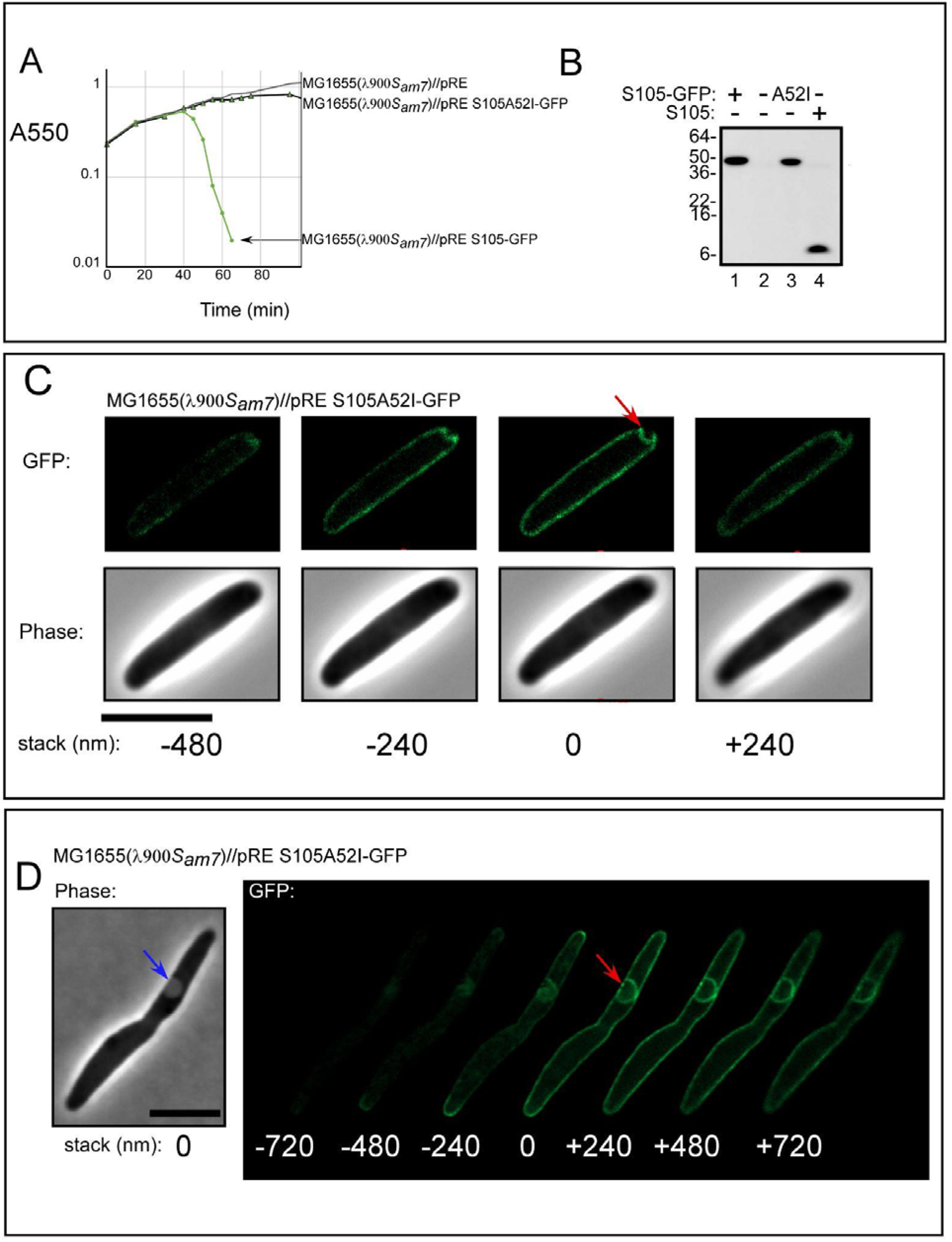
Rafts produced by lysis-defective S105A52I-GFP. λ900*S_am7_* lysogens induced for lysis and expressing S105A52I-GFP, isogenic to Fig. 5. A) Lysis profile showing complementation of *S_am7_* by plasmid-expressed *S105-GFP*. λ900*S_am7_*lysogens expressing pRE plasmids as indicated. B) Anti-S105 western blot comparing the accumulation of S105A52I-GFP to S105-GFP and S105. Representative polar (C) and non-polar (D) Deconvolved Z-stack images of λ900S*_am7_* lysogens induced for *S105_A52I_-GFP*. Cells were imaged 60 min after induction. Stack height is shown in nm, with “0” assigned to the in-focus frame. Red arrows indicate rafts. Blue arrows indicate phase-light spots. Scale bar = 5 µm

We tested S105_A52I_ using the same fixation and imaging conditions above with λΔ*SR* lysogens carrying an inducible plasmid, p*S105_A52I_*, that supplies the holin gene product in trans. Like the GFP-tagged construct, this mutant formed large rafts (Fig. S2B). Also like above, the A52I mutant had fewer rafts on average per cell (1.8 ± 0.92 rafts) compared to the wild-type holin strain.

To investigate these structures in three dimensions, we turned to imaging on a 3D SIM microscope (Materials and Methods, subsection Structured Illumination Microscopy (SIM) and analysis). As seen with z-stack imaging of S105_A52I-_GFP, this mutant formed large structures inside the cell that appeared to be present even at the mid-cell plane (Fig. S2C). Like the GFP experiments above, this suggests that the “ring” structures we observed in the superresolution images were actually three-dimensional rafts that had invaginated inside the cell cytoplasm. This also suggests that these raft structures invaginate the IM several hundred nanometers. Most cells imaged had rafts that localized both to the membrane z-plane as well as the cell mid-plane (66.67%, 42/63 cells). Additionally, consistent with the above results, the vast majority of cells had raft localizations at the poles of the cell (85%, 53/63 cells). Overall, the ImmunoSTORM and GFP data indicate that the rafts form most often at the poles and rafts predict the site of lysis (Table S2). Taken together, these results support the notion that the holin controls the site of lysis.

### Thioflavin-T labeling indicates IM permeabilization before lytic blowout

It was shown previously that loss of PMF preceded lysis by ∼ 19 ± 6 sec (see introduction). Moreover, the interval of time between PMF loss and lysis could be shortened by increasing the amount of endolysin [9], indicating that during a significant portion of this interval, the S- holes are large enough to release endolysin, presumably in the pathway to the formation of the micron-scale holes. To address this idea, a sensitive reporter system for IM permeabilization was developed. Thioflavin-T (ThT) has been shown to fluoresce upon binding to *E. coli* RNA and DNA [26]. ThT is soluble in water and small enough to enter the periplasm; therefore, we would expect ThT to serve as a reporter of IM permeabilization. To test this, we added ThT to log-phase cultures at the time of thermal induction. Cells were imaged prior to lysis using phase contrast and CFP filter settings. We monitored lysis of 42 cells and detected a continuous increase in ThT fluorescence before lysis (Fig. 7, Movie 9). The average time from increase in signal above background to lytic blowout was 13 seconds for λ900 (n =27, standard deviation = 5.3). The average time from ThT signal to the start of shape conversion for λ900*Rz_am_/Rz1_am_* lysogens was 10.3 seconds (Movie 10) (n = 15, standard deviation = 5.2).

**Figure 7.**
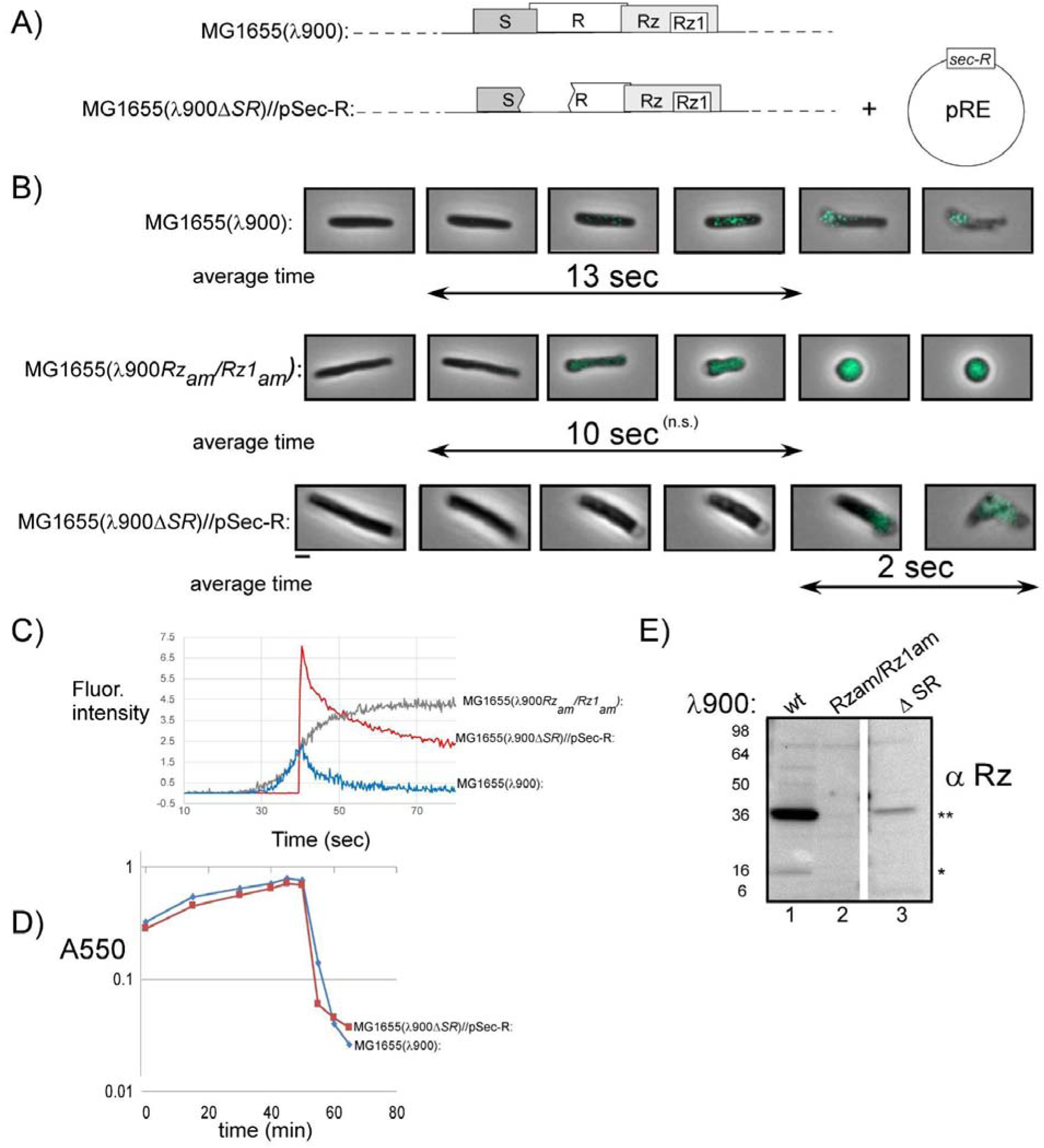
Thioflavin-T indicates inner membrane permeabilization before lysis. A) The genotypes used in the panels below. B) Thioflavin-T (ThT) was added to lysogens at the time of thermal induction. Cells were monitored 1 min prior to lysis with phase-contrast and CFP time-lapse microscopy. Micrographs are a merger of phase and CFP channels and show individual cells progressing through lysis or morphological conversion. Double arrows: average time from detection ThT signal above background to lysis or shape change is displayed below micrographs. Scale bar = 2 µm. C) Maximum fluorescence pixel intensity within cells plotted over time. Signals were normalized. Arbitrary units are shown for the y-axis. D) Lysis profile demonstrating the lytic phenotype of λ900Δ*SR* lysogens complemented by *Sec-R (phoA-R)* expression. Lysogens were induced at time 0 and monitored at A550. E) Western blots comparing Rz expressed from λ900 (wt), λ900*Rz_am_/Rz1_am_*, and λ900Δ*SR* lysogens. The position of the Rz monomer and covalently linked Rz dimer are indicated by single and double asterisks, respectively

To test whether ThT signal was dependent on IM permeabilization, we used lysogens encoding a deletion of the holin and endolysin (λ900Δ*SR*). In this background, the IM remains intact, since there is no holin activity. As expected, these cells showed no change in fluorescence during the same ThT treatment (not shown). This indicates that during lysis, ThT is labeling the cytoplasm as the IM becomes permeabilized by the holin. Therefore, if lysis could be induced independent of holin activity, we would expect ThT labeling to occur upon, but not before, breaching of the membrane. To address this, we used MG1655 cells expressing *^ss^PhoA-R* to complement the lysis defect of λ900(Δ*SR*) lysogens. As expected, we did not detect any fluorescence prior to lysis (Fig. 7B). Upon lytic blowout, the ThT signal was transiently detected, diffusing along with released cytoplasmic content (Fig. 7C) (Movie 11). Notably, these cells exhibited a rounding phenotype before lysis despite the lysis kinetics appearing normal in bulk culture (Fig. 7D). We attribute this defect to reduced spanin expression caused by the Δ*SR* deletion ∼100 bp upstream of *Rz* (Fig. 7E). These data indicate that ThT-permissive membrane S-holes form ∼10 seconds before lysis in λ lysogens.

## DISCUSSION

The first attempts to systematically document morphological changes during phage lysis can be traced to 1933 when photomicrographic analysis was used to capture lysis of *E. coli* at frame rates as fast as 8 per second [27]. Thus, the molecular basis of the morphological changes that occur during phage lysis have been a long-standing question. Recently, explosive cell lysis was reported to be crucial for biofilm development in *Pseudomonas* [28]. Super-resolution microscopy was used to show that lysis of a sub-population of cells within a biofilm produced vesicles and eDNA, which could be used by other cells. Therefore, the study of what happens to cells during phage lysis may have implications for human health. Phage λ is the most well-studied lysis system. Although previous reports have documented that lambda exhibits polar lysis morphology [11], the molecular basis for this was unclear. In this study we used fluorescence and phase time-lapse microscopy to track the distribution and rearrangement of lysis proteins in the seconds preceding lysis.

### Testing whether the endolysin causes the polar lysis phenotype

We tested the possible role of the endolysin in the polar lysis phenotype by inducing a chimera of the lambda endolysin (R) that was fused to the *phoA* signal sequence so R would be secreted into the periplasm in the absence of other lysis proteins. The uniform conversion to a spherical shape (Fig. 2C) is in contrast to that of λ900*Rz_am_/Rz1_am_* lysogens, in which R is released through S holes. At the time of lysis, such cells lose shape starting from the poles (Fig. 2B). We interpret this as evidence for the central role of the holin in affecting a polar lysis phenotype.

The finding that the endolysin is not biased to polar PG substrate is consistent with an *in vitro* study that demonstrated a homogenous susceptibility of purified *E. coli* sacculi to muramidase degradation [29].

### Holin holes

Almost two decades ago the minimum size of the S-hole was interrogated using a hybrid endolysin-ß-gal fusion, a 480 kDa homo-tetramer. Despite being more than 30-fold larger than the native R endolysin, lysis was indistinguishable both in kinetics and extent with this chimera [30]. Although this showed that S-holes were large enough to permit the transit of the chimeric endolysin across the IM, the actual hole size was not determined until cryoEM methods were used. Later, cysteine-scanning accessibility showed that virtually all molecules of S105 participate in hole formation and that the hydrophilic faces of two S105 TMDs face the lumen of the hole [31]. The number of S105 molecules per cell was determined by quantitative western blotting to be ∼1000 [32] and each TMD occupies ∼1 nm of area. Therefore, 1000 copies of S105 would produce 2000 nm of hole-lining perimeter. This would account for a single 600 nm-diameter hole or two 300 nm holes, which is consistent with the number and diameter of holes observed by cryoEM of S105-expressing cells [5, 7, 8]. CryoEM showed that hole size ranged from ∼100-1000 nm in diameter, averaging 340 nm. Up to four holes per cell were detected, and appeared to be randomly positioned in the IM. Estimates suggest there are two holes per cell on average when taking factors such as geometry of viewing, observed diameters and locations of holes into account. Notably, cryoEM could not be used to address the question of polar lysis since the cell poles were not included in the analysis. This was because the IM appeared discontinuous at the poles of control cells that were not expressing the holin [8]. Generally, most of the recent studies have focused on the endpoint of holin function and have not been able to associate the dynamics of the holin and the relation of rafts to lysis morphology in lysing cells.

### Model for holin function

Based on the findings above, we incorporate our observations of S105-GFP rearrangement into the current model of holin function (Fig. 8). To account for the evidence of large hole formation and precise timing of the holin, it is thought that the “all-or-nothing” response is set to an allele-specific critical concentration (Fig. 8A). At this point the holin undergoes a dramatic transition to form ∼2 rafts per cell about 100 sec prior to lysis (Fig. 4B, Table S2, Fig. 8). The critical concentration-dependent transition for membrane proteins has precedent in halobacteria, where bacteriorhodopsin (BR) forms a two-dimensional lattice (purple membrane) [33]. Unlike BR, holin rafts are thought to cause a collapse of membrane potential, at which time the rafts rearrange into hole-forming structures that initiate lysis. Previous studies using an assay based on flagella rotation showed that PMF loss occurs ∼19 ±6 seconds [9]. Here, we used ThT to report on cell permeabilization based on the increase in fluorescence of ThT upon entering the cytoplasm (Fig. 7). ThT labeling occurs ∼13 ±5 seconds before lysis, indicating that the hole size has increased large enough to permit ThT entry (Fig. 8A). Our data indicates that the rafts form most often at the poles, causing the subsequent steps of lysis (hole formation, endolysin release) to also occur at the poles. Spanins disrupt the OM at the site of PG degradation, completing the last step of the lysis pathway. Taken together, the data indicate that the holin controls the site of lysis and directs lysis from the poles.

**Figure 8.**
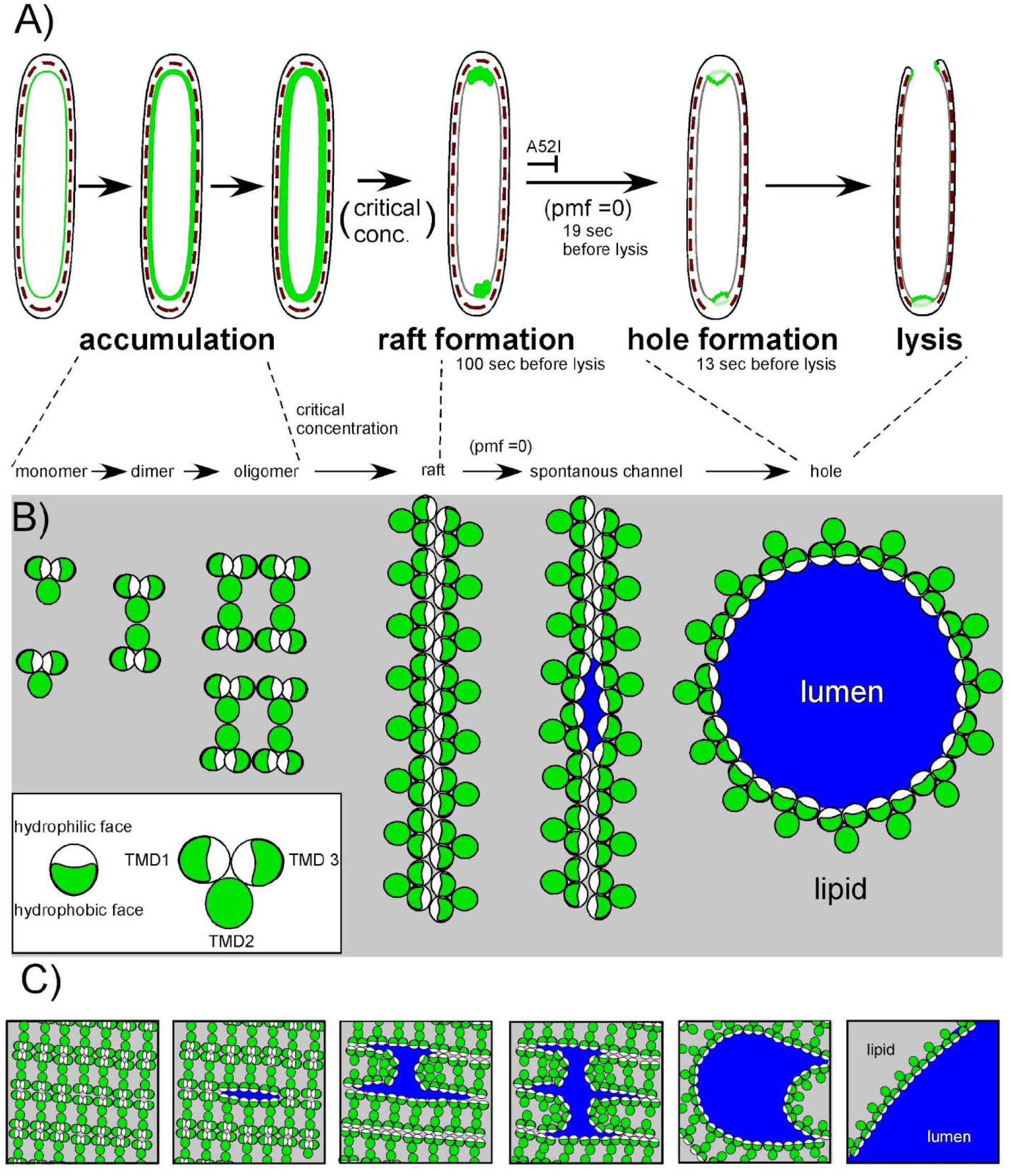
Model of holin function incorporated into observed morphological changes prior to lysis. A) The holin is shown in green accumulating in the IM and nucleates into rafts upon reaching a critical concentration. Rafts are most often associated with the poles. Rafts trigger, forming holes in the IM which result in localized blowout proximal to the site of raft formation. B) Top-down view of the holin. As the holin accumulates in the IM, the hydrophilic faces (white) of TMD1 and 3 are in contact, sequestered from the lipid. In the raft form the hydrophilic faces are associated by intermolecular contacts. Rafts persist until the energy production of the cell is blocked. When pmf is lost, the raft rearranges into a hole-forming structure, which allows endolysin access to the periplasm. Lysis begins. For the *S105_A52I_*product, holin function is stalled after forming the raft superstructure and do not form holes. The cartoons of the rafts and the hole are not drawn to scale. The linear arrangement has not been demonstrated and is shown purely to indicate the sequestration of the hydrophilic faces of the holin molecules from the lipid. C) Model for how a holin raft array transitions to the hole- forming arrangement.

### The holin “death rafts”

A key remaining question is how do rafts lead to holes? Nothing is known about the organization of holin molecules in the raft structure or about how rafts transition to hole-forming arrangements. In contrast to the hypothetical linear structures in Fig. 8B the GFP data demonstrate that rafts were much more densely packed. Thus, the rafts are more likely to instead be composed of repeated 2D linear arrangements juxtaposed in an array-like structure (Fig. 8C). To address the question of how rafts disrupt the PMF, our lab has proposed the “death raft” model, in which the intimate packing of the holin creates a protein-rich, lipid-depleted area that is a poor insulator and causes a local collapse in PMF [6]. The array arrangement partially solves the problem of how to remove lipids from a hole because the interior of the raft array may be so densely packed that it is already significantly lipid-depleted. However, without further structural data, it is unclear how a large hole(s) forms from the array structure. Nevertheless, we present a simple model for array-hole transition, supposing that upon PMF collapse the holin hydrophilic TMD contacts suddenly reorient to permit the expansion of a large hole (Fig. 8C).

Lastly, in this report we present evidence that S105_A52I_ is blocked at the hole formation step and this allele might be used to determine the orientation of holin molecules within the raft. Future experiments, perhaps using correlative cryoEM and molecular modeling could address the structure of rafts and probe the raft-to-hole transition during lysis.

---

**Figure S1.**
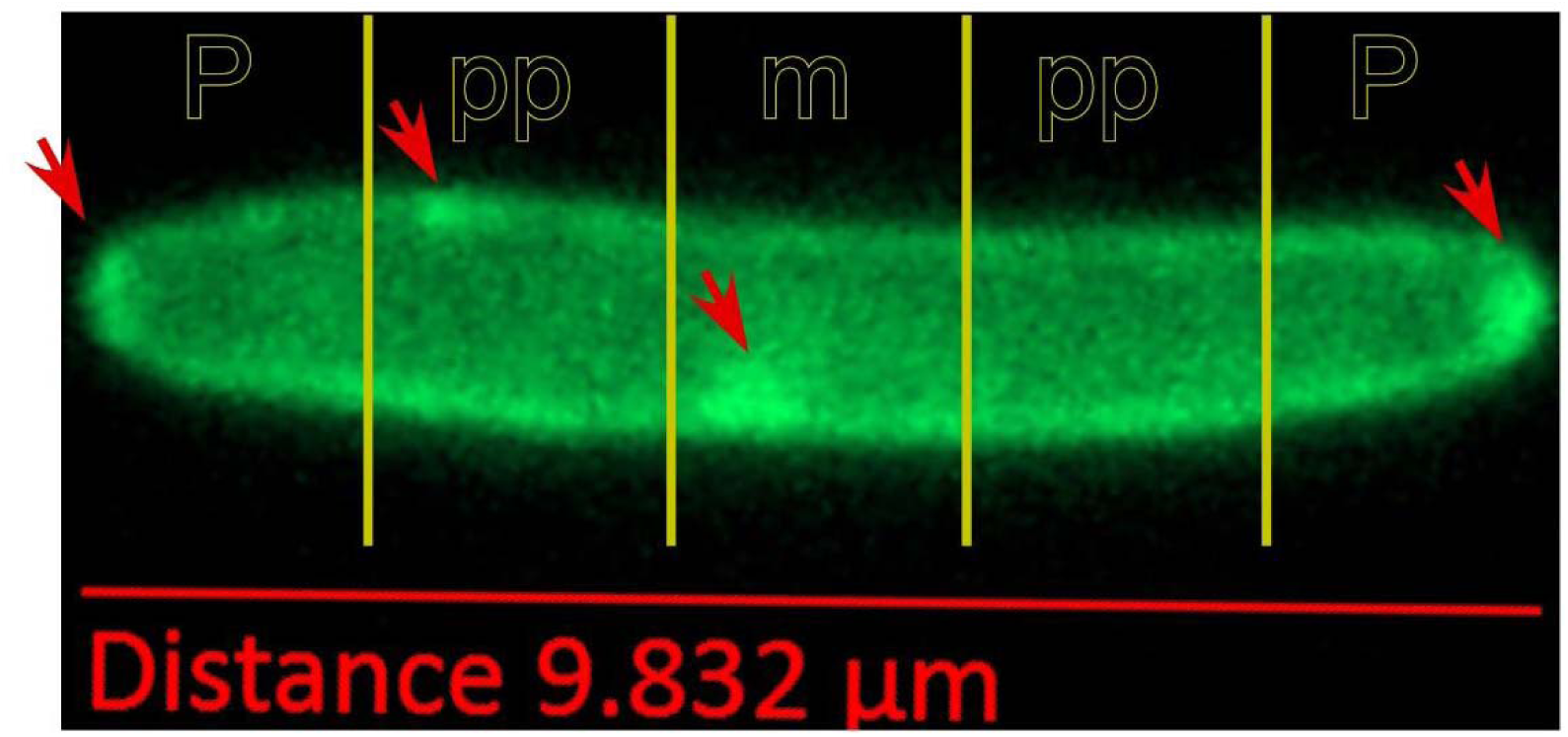
Assigning rafts to subcellular compartments. P=polar, pp=peripolar, m=midcell.

**Figure S2.**
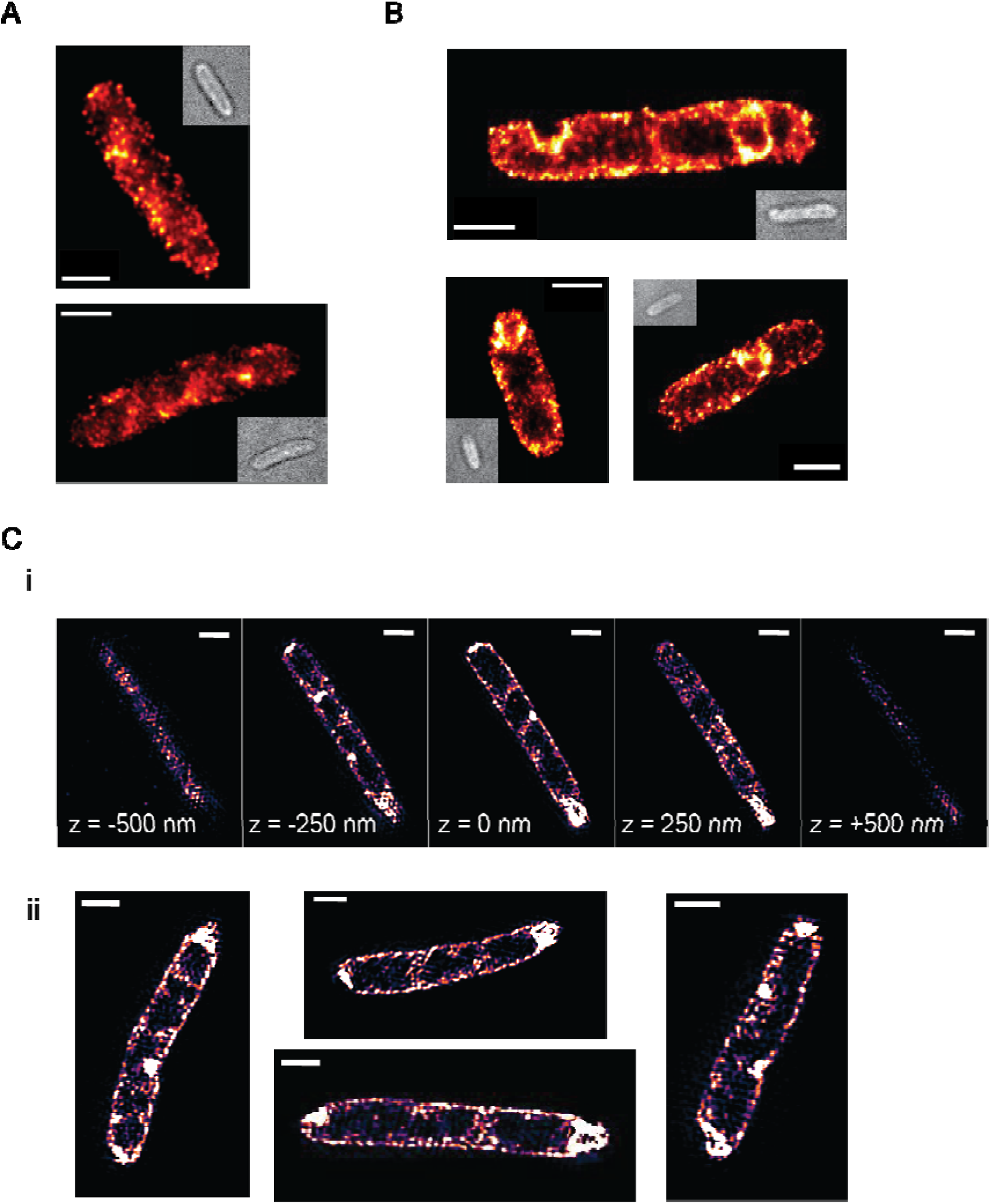
Immuno-STORM imaging of S105 and S105-A52I. Superresolution imaging of S105 and the A52I non-lytic holin mutant. Representative STORM images of (A) WT holin at 50 min post-induction of MG1655(λ*R_am_*), and (B) λ900 ΔSR lysogens induced for *S105_A52I_* imaged at 50 min post-induction, inset images are brightfield images of the cells; each superresolution image contains fitted spots represented as two- dimensional Gaussians, normalized against the total intensity, scale bars represent one micron. (C) Reconstructed SIM images of the A52I non-lytic holin mutant at 50 min post-induction, with (i) showing a montage of a single cell in the z-dimension demonstrating an invaginated holin raft with (ii) showing representative images of rafts at the mid-cell plane; scale bars represent one micron.

**Table S1.**
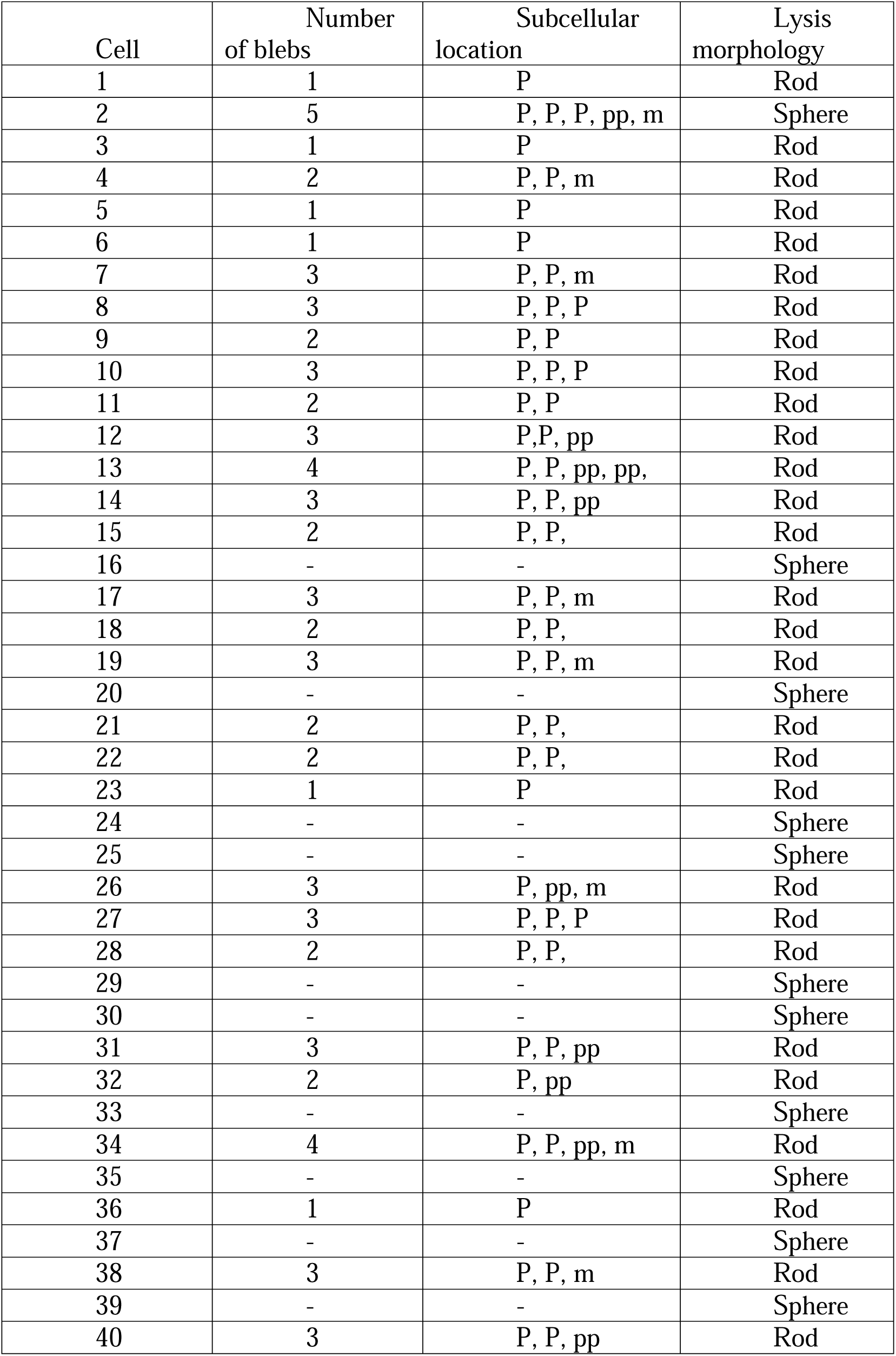
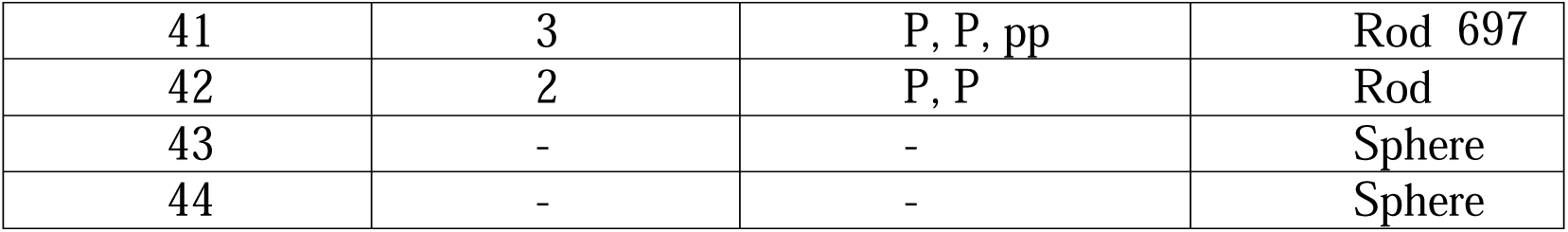
Lysis of spanin mutant cells treated with EDTA prior to lysis. The number of blebs detected per cell, subcellular localization of blebs and lysis morphology are described per cell. P = polar; pp = parapolar; m = midcell. “-” indicates blebs were not detected.

**Table S2.**
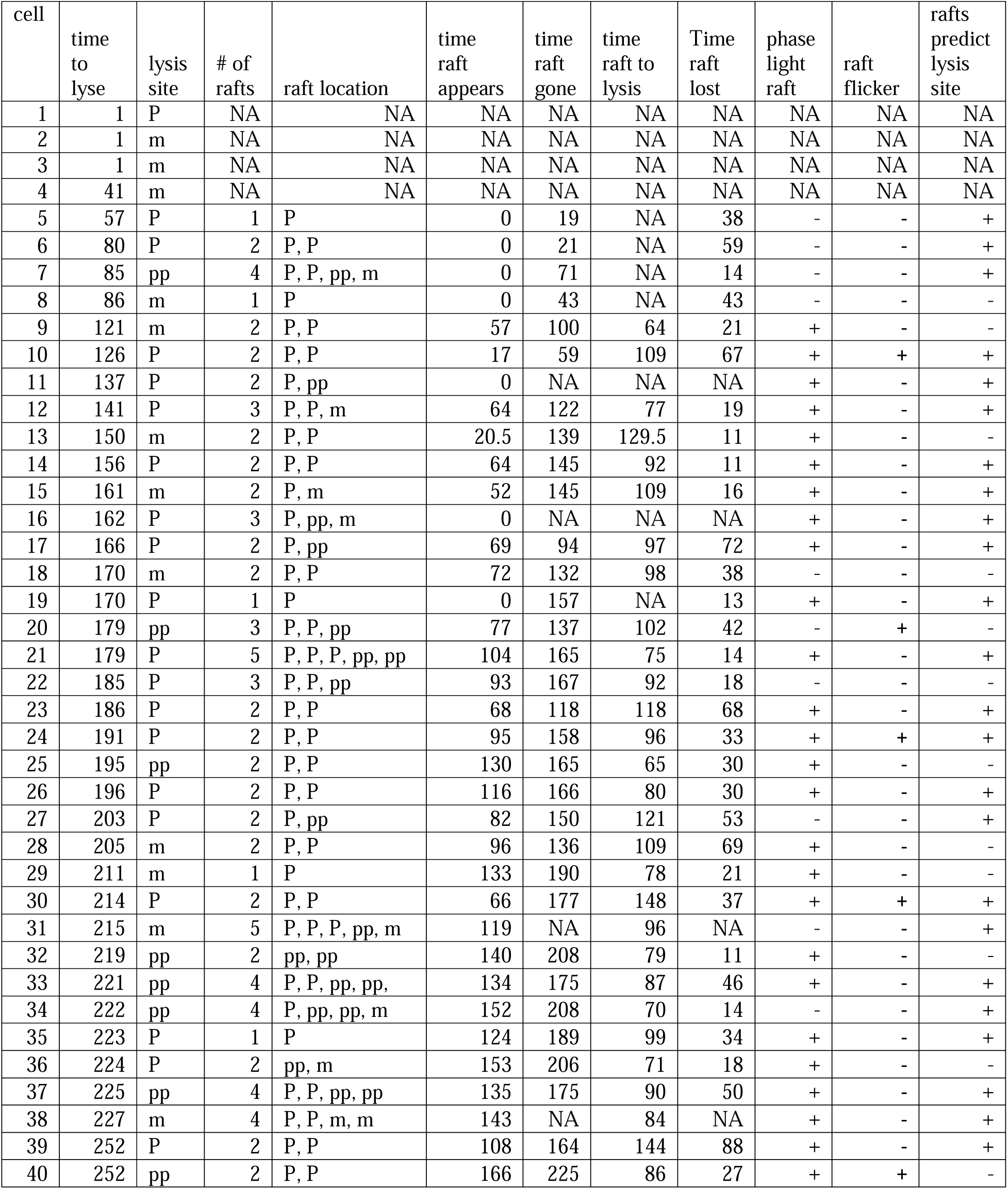
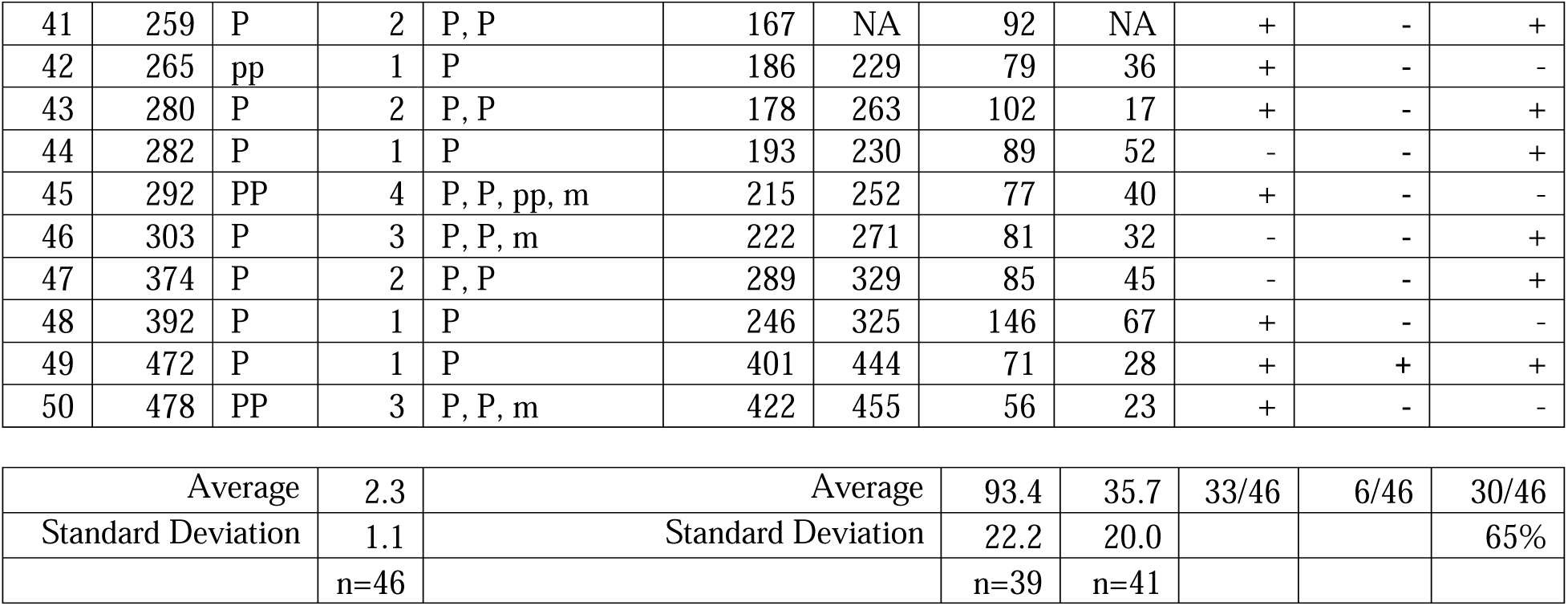
Lysis of cells expressing *S105-GFP.* The table is ordered by the monitoring time before lysis. Time is reported in seconds. Breach site and raft locations are indicated by P=polar, m= midcell, pp =parapolar. NA. = uncertain because the parameter could not be detected. The “time raft to lysis” measures the seconds between raft formation and lysis. “time raft lost” is the interval of time from raft disappearance to lysis.

## Notes

### Competing Interest Statement

The authors have declared no competing interest.

## WORKS CITED

[1] R. Young, “Phage lysis: do we have the hole story yet?,” (in Eng), Curr Opin Microbiol, Oct 7 2013, doi: S1369-5274(13)00150-1 [pii] 10.1016/j.mib.2013.08.008.

[2] I. N. Wang, D. L. Smith, and R. Young, “Holins: the protein clocks of bacteriophage infections,” Annu. Rev. Microbiol., vol. 54, pp. 799–825, 2000. [Online]. Available: PM:0011018145

[3] R. Young, “Bacteriophage lysis: mechanism and regulation,” Microbiol. Rev., vol. 56, pp. 430–481, 1992.

[4] A. Gründling, D. L. Smith, U. Bläsi, and R. Young, “Dimerization between the holin and holin inhibitor of phage lambda,” J. Bacteriol., vol. 182, no. 21, pp. 6075–6081, 2000. [Online]. Available: PM:0011029427

[5] R. White et al., “Holin triggering in real time,” (in eng), Proc Natl Acad Sci U S A, vol. 108, no. 2, pp. 798–803, Jan 11 2011, doi: 1011921108 [pii] 10.1073/pnas.1011921108.

[6] J. Cahill and R. Young, “Phage lysis: multiple genes for multiple barriers,” Advances in virus research, vol. 103, pp. 33–70, 2019.

[7] J. S. Dewey, C. G. Savva, R. L. White, S. Vitha, A. Holzenburg, and R. Young, “Micron-scale holes terminate the phage infection cycle,” (in Eng), Proc Natl Acad Sci U S A, vol. 107, pp. 2219–2223, Jan 11 2010, doi: 0914030107 [pii] 10.1073/pnas.0914030107.

[8] C. G. Savva, J. S. Dewey, S. H. Moussa, K. H. To, A. Holzenburg, and R. Young, “Stable micron-scale holes are a general feature of canonical holins,” Molec Microbiol, vol. 91, no. 1, pp. 57–65, Nov 21 2014, doi: 10.1111/mmi.12439.

[9] A. Gründling, M. D. Manson, and R. Young, “Holins kill without warning,” Proc. Natl. Acad. Sci. U S A, vol. 98, no. 16, pp. 9348–9352, 2001. [Online]. Available: PM:11459934

[10] U. Bläsi, C. Y. Chang, M. T. Zagotta, K. Nam, and R. Young, “The lethal λ *S* gene encodes its own inhibitor,” EMBO Journal, vol. 9, pp. 981–989, 1990.

[11] J. Berry, M. Rajaure, T. Pang, and R. Young, “The spanin complex is essential for lambda lysis,” J. Bacteriol., vol. 194, pp. 5667–5674, 2012. [Online]. Available: http://www.ncbi.nlm.nih.gov/pubmed/22904283.

[12] J. Berry, E. J. Summer, D. K. Struck, and R. Young, “The final step in the phage infection cycle: the Rz and Rz1 lysis proteins link the inner and outer membranes, (in eng), Molec Microbiol, vol. 70, no. 2, pp. 341–51, Oct 2008, doi: MMI6408 [pii] 10.1111/j.1365-2958.2008.06408.x.

[13] J. Cahill et al., “Genetic analysis of the lambda spanins Rz and Rz1: identification of functional domains,” G3: Genes| Genomes| Genetics, p. g3. 116.037192, 2016.

[14] N. Zhang and R. Young, “Complementation and characterization of the nested *Rz* and *Rz1* reading frames in the genome of bacteriophage lambda,” Mol. Gen. Genet., vol. 262, no. 4-5, pp. 659–667, 1999. [Online]. Available: PM:0010628848

[15] M. Rajaure, J. Berry, R. Kongari, J. Cahill, and R. Young, “Membrane fusion during phage lysis, (in eng), Proceedings of the National Academy of Sciences of the United States of America, Research Support, N.I.H., Extramural Research Support, Non-U.S. Gov’t vol. 112, no. 17, pp. 5497–502, Apr 28 2015, doi: 10.1073/pnas.1420588112.

[16] A. Gründling, U. Bläsi, and R. Young, “Biochemical and genetic evidence for three transmembrane domains in the class I holin, lambda S,” (in eng), J Biol Chem, vol. 275, no. 2, pp. 769–76, Jan 14 2000. [Online]. Available: http://www.ncbi.nlm.nih.gov/entrez/query.fcgi?cmd=Retrieve&db=PubMed&dopt=Citation&list_uids=10625606.

[17] J. Cahill et al., “Suppressor analysis of the fusogenic lambda spanins,” Journal of virology, pp. JVI. 00413-17, 2017.

[18] L. Leive, “The barrier function of the gram-negative envelope,” Annals of the New York Academy of Sciences, vol. 235, no. 0, pp. 109–29, May 10 1974. [Online]. Available: http://www.ncbi.nlm.nih.gov/pubmed/4212391.

[19] D. L. Smith, D. K. Struck, J. M. Scholtz, and R. Young, “Purification and biochemical characterization of the lambda holin,” J. Bacteriol., vol. 180, no. 9, pp. 2531–2540, 1998.

[20] R. White, T. A. Tran, C. A. Dankenbring, J. Deaton, and R. Young, “The N-terminal transmembrane domain of lambda S is required for holin but not antiholin function,” (in eng), Journal of bacteriology, Research Support, N.I.H., Extramural Research Support, Non-U.S. Gov’t vol. 192, no. 3, pp. 725–33, Feb 2010, doi: 10.1128/JB.01263-09.

[21] C. Paradis-Bleau et al., “Lipoprotein cofactors located in the outer membrane activate bacterial cell wall polymerases,” Cell, vol. 143, no. 7, pp. 1110–1120, 2010.

[22] R. Johnson-Boaz, C. Y. Chang, and R. Young, “A dominant mutation in the bacteriophage lambda *S* gene causes premature lysis and an absolute defective plating phenotype,” Mol. Microbiol., vol. 13, pp. 495–504, 1994.

[23] R. Raab, G. Neal, J. Garrett, R. Grimaila, R. Fusselman, and R. Young, “Mutational analysis of bacteriophage lambda lysis gene *S*” J.Bacteriology, vol. 167, pp. 1035–1042, 1986.

[24] R. Raab, G. Neal, C. Sohaskey, J. Smith, and R. Young, “Dominance in lambda *S* mutations and evidence for translational control,” J. Mol. Biol., vol. 199, pp. 95–105, 1988.

[25] R. L. White, “What Makes the Lysis Clock Tick?: A Study of the Bacteriophage Lambda Holin,” Texas A & M University, 2010.

[26] S. Sugimoto, K.-i. Arita-Morioka, Y. Mizunoe, K. Yamanaka, and T. Ogura, “Thioflavin T as a fluorescence probe for monitoring RNA metabolism at molecular and cellular levels,” Nucleic Acids Res., vol. 43, no. 14, pp. e92–e92, 2015.

[27] S. Bayne-Jones and L. A. Sandholzer, “Changes in the shape and size of Bacterium coli and Bacillus megatherium under the influence of bacteriophage—A motion photomicrographic analysis of the mechanism of lysis,” Journal of Experimental Medicine, vol. 57, no. 2, pp. 279–303, 1933.

[28] L. Turnbull et al., “Explosive cell lysis as a mechanism for the biogenesis of bacterial membrane vesicles and biofilms,” Nature communications, vol. 7, 2016.

[29] M. A. de Pedro, J. C. Quintela, J. V. H”ltje, and H. Schwarz, “Murein segregation in Escherichia coli ” J.Bacteriology, vol. 179, no. 9, pp. 2823–2834, 1997. [Online]. Available: PM:0009139895

[30] I.-N. Wang, J. Deaton, and R. Young, “Sizing the Holin Lesion with an Endolysin-β- Galactosidase Fusion,” J. Bacteriol., vol. 185, no. 3, pp. 779–787, February 1, 2003 2003, doi: 10.1128/jb.185.3.779-787.2003.

[31] K. H. To and R. Young, “Probing the structure of the S105 hole,” (in Eng), J Bacteriol, Aug 4 2014, doi: 10.1128/jb.01673-14.

[32] C. Y. Chang, K. Nam, and R. Young, “S gene expression and the timing of lysis by bacteriophage lambda,” (in eng), J Bacteriol, vol. 177, no. 11, pp. 3283–94, Jun 1995. [Online]. Available: http://www.ncbi.nlm.nih.gov/entrez/query.fcgi?cmd=Retrieve&db=PubMed&dopt=Citation&list_uids=7768829.

[33] M. P. Krebs and T. A. Isenbarger, “Structural determinants of purple membrane assembly,” Biochimica et Biophysica Acta (BBA)-Bioenergetics, vol. 1460, no. 1, pp. 15–26, 2000.

