## supplementary tables for "Spatial and temporal control of lysis by the lambda holin"

**Table S1. Lysis of spanin mutant cells treated with EDTA prior to lysis. The number of blebs detected per cell, subcellular localization of blebs and lysis morphology are described per cell. P = polar; pp = parapolar; m = midcell. “-” indicates blebs were not detected.**

| Cell | Number of blebs | Subcellular location | Lysis morphology |
| --- | --- | --- | --- |
| 1 | 1 | P | Rod |
| 2 | 5 | P, P, P, pp, m | Sphere |
| 3 | 1 | P | Rod |
| 4 | 2 | P, P, m | Rod |
| 5 | 1 | P | Rod |
| 6 | 1 | P | Rod |
| 7 | 3 | P, P, m | Rod |
| 8 | 3 | P, P, P | Rod |
| 9 | 2 | P, P | Rod |
| 10 | 3 | P, P, P | Rod |
| 11 | 2 | P, P | Rod |
| 12 | 3 | P,P, pp | Rod |
| 13 | 4 | P, P, pp, pp, | Rod |
| 14 | 3 | P, P, pp | Rod |
| 15 | 2 | P, P, | Rod |
| 16 | - | - | Sphere |
| 17 | 3 | P, P, m | Rod |
| 18 | 2 | P, P, | Rod |
| 19 | 3 | P, P, m | Rod |
| 20 | - | - | Sphere |
| 21 | 2 | P, P, | Rod |
| 22 | 2 | P, P, | Rod |
| 23 | 1 | P | Rod |
| 24 | - | - | Sphere |
| 25 | - | - | Sphere |
| 26 | 3 | P, pp, m | Rod |
| 27 | 3 | P, P, P | Rod |
| 28 | 2 | P, P, | Rod |
| 29 | - | - | Sphere |
| 30 | - | - | Sphere |
| 31 | 3 | P, P, pp | Rod |
| 32 | 2 | P, pp | Rod |
| 33 | - | - | Sphere |
| 34 | 4 | P, P, pp, m | Rod |
| 35 | - | - | Sphere |
| 36 | 1 | P | Rod |
| 37 | - | - | Sphere |
| 38 | 3 | P, P, m | Rod |
| 39 | - | - | Sphere |
| 40 | 3 | P, P, pp | Rod |
| 41 | 3 | P, P, pp | Rod |
| 42 | 2 | P, P | Rod |
| 43 | - | - | Sphere |
| 44 | - | - | Sphere |

**Table S2. Lysis of cells expressing *S105-GFP.* The table is ordered by the monitoring time before lysis. Time is reported in seconds. Breach site and raft locations are indicated by P=polar, m= midcell, pp =parapolar. NA. = uncertain because the parameter could not be detected. The “time raft to lysis” measures the seconds between raft formation and lysis. “time raft lost” is the interval of time from raft disappearance to lysis.**

| cell | time to lyse | lysis site | # of rafts | raft location | time raft appears | time raft gone | time raft to lysis | | Time raft lost | | phase light raft | | raft flicker | | rafts predict  lysis site | |
| --- | --- | --- | --- | --- | --- | --- | --- | --- | --- | --- | --- | --- | --- | --- | --- | --- |
| 1 | 1 | P | NA | NA | NA | NA | NA | | NA | | NA | | NA | | NA | |
| 2 | 1 | m | NA | NA | NA | NA | NA | | NA | | NA | | NA | | NA | |
| 3 | 1 | m | NA | NA | NA | NA | NA | | NA | | NA | | NA | | NA | |
| 4 | 41 | m | NA | NA | NA | NA | NA | | NA | | NA | | NA | | NA | |
| 5 | 57 | P | 1 | P | 0 | 19 | NA | | 38 | | - | | - | | + | |
| 6 | 80 | P | 2 | P, P | 0 | 21 | NA | | 59 | | - | | - | | + | |
| 7 | 85 | pp | 4 | P, P, pp, m | 0 | 71 | NA | | 14 | | - | | - | | + | |
| 8 | 86 | m | 1 | P | 0 | 43 | NA | | 43 | | - | | - | | - | |
| 9 | 121 | m | 2 | P, P | 57 | 100 | 64 | | 21 | | + | | - | | - | |
| 10 | 126 | P | 2 | P, P | 17 | 59 | 109 | | 67 | | + | | + | | + | |
| 11 | 137 | P | 2 | P, pp | 0 | NA | NA | | NA | | + | | - | | + | |
| 12 | 141 | P | 3 | P, P, m | 64 | 122 | 77 | | 19 | | + | | - | | + | |
| 13 | 150 | m | 2 | P, P | 20.5 | 139 | 129.5 | | 11 | | + | | - | | - | |
| 14 | 156 | P | 2 | P, P | 64 | 145 | 92 | | 11 | | + | | - | | + | |
| 15 | 161 | m | 2 | P, m | 52 | 145 | 109 | | 16 | | + | | - | | + | |
| 16 | 162 | P | 3 | P, pp, m | 0 | NA | NA | | NA | | + | | - | | + | |
| 17 | 166 | P | 2 | P, pp | 69 | 94 | 97 | | 72 | | + | | - | | + | |
| 18 | 170 | m | 2 | P, P | 72 | 132 | 98 | | 38 | | - | | - | | - | |
| 19 | 170 | P | 1 | P | 0 | 157 | NA | | 13 | | + | | - | | + | |
| 20 | 179 | pp | 3 | P, P, pp | 77 | 137 | 102 | | 42 | | - | | + | | - | |
| 21 | 179 | P | 5 | P, P, P, pp, pp | 104 | 165 | 75 | | 14 | | + | | - | | + | |
| 22 | 185 | P | 3 | P, P, pp | 93 | 167 | 92 | | 18 | | - | | - | | - | |
| 23 | 186 | P | 2 | P, P | 68 | 118 | 118 | | 68 | | + | | - | | + | |
| 24 | 191 | P | 2 | P, P | 95 | 158 | 96 | | 33 | | + | | + | | + | |
| 25 | 195 | pp | 2 | P, P | 130 | 165 | 65 | | 30 | | + | | - | | - | |
| 26 | 196 | P | 2 | P, P | 116 | 166 | 80 | | 30 | | + | | - | | + | |
| 27 | 203 | P | 2 | P, pp | 82 | 150 | 121 | | 53 | | - | | - | | + | |
| 28 | 205 | m | 2 | P, P | 96 | 136 | 109 | | 69 | | + | | - | | - | |
| 29 | 211 | m | 1 | P | 133 | 190 | 78 | | 21 | | + | | - | | - | |
| 30 | 214 | P | 2 | P, P | 66 | 177 | 148 | | 37 | | + | | + | | + | |
| 31 | 215 | m | 5 | P, P, P, pp, m | 119 | NA | 96 | | NA | | - | | - | | + | |
| 32 | 219 | pp | 2 | pp, pp | 140 | 208 | 79 | | 11 | | + | | - | | - | |
| 33 | 221 | pp | 4 | P, P, pp, pp, | 134 | 175 | 87 | | 46 | | + | | - | | + | |
| 34 | 222 | pp | 4 | P, pp, pp, m | 152 | 208 | 70 | | 14 | | - | | - | | + | |
| 35 | 223 | P | 1 | P | 124 | 189 | 99 | | 34 | | + | | - | | + | |
| 36 | 224 | P | 2 | pp, m | 153 | 206 | 71 | | 18 | | + | | - | | - | |
| 37 | 225 | pp | 4 | P, P, pp, pp | 135 | 175 | 90 | | 50 | | + | | - | | + | |
| 38 | 227 | m | 4 | P, P, m, m | 143 | NA | 84 | | NA | | + | | - | | + | |
| 39 | 252 | P | 2 | P, P | 108 | 164 | 144 | | 88 | | + | | - | | + | |
| 40 | 252 | pp | 2 | P, P | 166 | 225 | 86 | | 27 | | + | | + | | - | |
| 41 | 259 | P | 2 | P, P | 167 | NA | 92 | | NA | | + | | - | | + | |
| 42 | 265 | pp | 1 | P | 186 | 229 | 79 | | 36 | | + | | - | | - | |
| 43 | 280 | P | 2 | P, P | 178 | 263 | 102 | | 17 | | + | | - | | + | |
| 44 | 282 | P | 1 | P | 193 | 230 | 89 | | 52 | | - | | - | | + | |
| 45 | 292 | PP | 4 | P, P, pp, m | 215 | 252 | 77 | | 40 | | + | | - | | - | |
| 46 | 303 | P | 3 | P, P, m | 222 | 271 | 81 | | 32 | | - | | - | | + | |
| 47 | 374 | P | 2 | P, P | 289 | 329 | 85 | | 45 | | - | | - | | + | |
| 48 | 392 | P | 1 | P | 246 | 325 | 146 | | 67 | | + | | - | | - | |
| 49 | 472 | P | 1 | P | 401 | 444 | 71 | | 28 | | + | | + | | + | |
| 50 | 478 | PP | 3 | P, P, m | 422 | 455 | 56 | | 23 | | + | | - | | - | |
| Average | | | 2.3 | Average | | | | 93.4 | | 35.7 | | 33/46 | | 6/46 | | 30/46 |
| Standard Deviation | | | 1.1 | Standard Deviation | | | | 22.2 | | 20.0 | |  | |  | | 65% |
|  | | | n=46 |  | | | | n=39 | | n=41 | |  | |  | |  |
